# The evidence for anthocyanins in the betalain-pigmented genus Hylocereus is weak

**DOI:** 10.1101/2021.11.16.468878

**Authors:** Boas Pucker, Samuel F. Brockington

## Abstract

Here we respond to Zhou et al., 2020 “Combined Transcriptome and Metabolome analysis of Pitaya fruit unveiled the mechanisms underlying Peel and pulp color formation” published in BMC Genomics. Given the evolutionary conserved anthocyanin biosynthesis pathway in betalain-pigmented species, we are open to the idea that species with both anthocyanins and betalains might exist. However, in absence of LC-MS/MS spectra, apparent lack of biological replicates, and no comparison to authentic standards, the findings of Zhou et al., 2020 are not a strong basis to propose the presence of anthocyanins in betalain-pigmented pitaya. In addition, our re-analysis of the datasets indicates the misidentification of important genes and the omission of key anthocyanin synthesis genes *ANS* and *DFR*. Finally, our re-analysis of the RNA-Seq dataset reveals no correlation between anthocyanin biosynthesis gene expression and pigment status.

## MAIN TEXT

Betalain pigments are restricted to the Caryophyllales, where they replace the otherwise ubiquitous anthocyanins in several families [1]. However, not all families in the Caryophyllales produce betalains. The complex pigment distribution over evolutionary lineages can be explained by at least four independent origins of the betalain biosynthesis [2]. Interestingly, anthocyanins have not been observed in betalain-pigmented species [1, 3, 4]. Consequently, the theory of mutual exclusion of both pigments was established and repeatedly supported by numerous studies in the last decades [3, 4]. Although co-occurrence of anthocyanins and betalains can be achieved through genetic engineering [5], simultaneous accumulation of both pigments within a native species has previously never been reported in nature. Zhou et al. 2020 [6] stated that “the anthocyanin coexistence with betalains is unneglectable” in their publication about the pigmentation of pitayas. Here, we outline some reasons why we do not think the study by Zhou et al.,2020 provides solid evidence for the presence of anthocyanins in betalain-pigmented pitayas.

### Patterns of proposed anthocyanin accumulation are weak basis for subsequent correlation with genes expression

Zhou et al., investigated metabolic differences between three pitaya cultivars: red peel/red pulp (RR), yellow peel/white pulp (YW), and green peel/white pulp (GW). They claim that 70 different anthocyanins are differentially accumulated in peels and pulps of these cultivars, but Table S9 lists only 14 anthocyanins. Unfortunately, these crucial metabolic analyses were apparently restricted to a single sample per cultivar (Table S9) which prevents any solid conclusions about the quantity of pigments. Nevertheless, we calculated the total amounts of detected anthocyanins per tissue and cultivar based on the data they provide (**Table 1**). While the total anthocyanin amount in the red peel is substantially higher than the amounts in yellow or green peel, there is only a very small difference between the different pulp samples. The difference between the two white pulps is substantially higher than the difference between one white and the red pulp. There are more anthocyanins reported in the yellow or green peel than in the red pulp. The detection of delphinidin and malvidin glycosides (blue pigments) would imply the presence of a functional flavonoid 3’,5’-hydroxylase (F3’5’H) in all three cultivars. However, Zhou et al., do not mention such an enzyme and our analyses revealed no evidence for the presence of *F3’5’H* transcripts in pitaya. Zhou et al., report that “metabolites with similar fragment ions were suggested to be the same compounds”. But as we previously outlined this method does not follow best practice [7]. We would expect at least to see the LC-MS/MS spectra and co-elution/fragmentation of the pigments versus authentic reference compounds [7], especially when reporting the unexpected occurrence of anthocyanins in a betalain-pigmented species.

**Table 1:** Total suggested anthocyanin amount reported by Zhou et al., 2020 in three pitaya cultivars.

| Cultivar/tissue | Total amount of suggested anthocyanins |
| --- | --- |
| RR-peel | 6,669,221 |
| GW-peel | 786,331 |
| YW-peel | 797,863 |
| RR-pulp | 631,184 |
| GW-pulp | 479,385 |
| YW-pulp | 155,051 |

### Transcriptomic analysis indicates likely block in anthocyanidin biosynthesis at the level of DFR and ANS genes

Zhou et al., claim “Our results demonstrated that anthocyanin biosynthesis was one of the significantly enriched pathways”. We do not see any evidence for this statement, because the analysis presented in their Fig. 5 covers only the general phenylpropanoid pathway and selected steps of the flavonoid biosynthesis. Our re-evaluation based on the construction of gene trees for all steps in the pathway (**Additional file 1, Additional file 2**) indicates several cases of misidentification and missed gene copies, although their annotation is difficult to evaluate as multiple transcripts are reported for each gene class, and clear orthology is not established in absence of a phylogenetic analysis. Important genes of the anthocyanin biosynthesis like *DFR* and *ANS* are missing. For example, *DFR* (Cluster-16519.0) and *ANS* (Cluster-7001.0) are both present in the transcriptome assembly, but were not presented by Zhou *et al*. Zhou et al., write “The comparative pitaya transcriptome showed the differential regulation of the anthocyanin pathway and genes controlling almost every single step in the pathway were differentially regulated”. However, two very important anthocyanin biosynthesis genes, *DFR* and *ANS*, are not presented in Fig. 5 of Zhou et al. Since expression of *DFR* and *ANS* would be crucial for the formation of anthocyanins, we performed a re-analysis of the flavonoid biosynthesis including genes for the reactions leading to anthocyanidins (**Fig. 1**). Our analysis is based on a transcriptome assembly of RR (red fruits) and indicates a block in the anthocyanin biosynthesis at DFR and ANS (**Fig. 1**). We would expect to see high transcript abundance of all genes necessary for anthocyanin formation if anthocyanins would be substantially contributing to the red colour, especially at the intensity they report. We included *LAR* and *ANR* in our analysis because the enzymes encoded by these genes are responsible for proanthocyanidin biosynthesis. Anthocyanins and proanthocyanins have shared precursors and could be considered as competing pathways. The presence of ANR transcript at high levels indicates that the lowly abundant DFR and ANS transcripts might be involved in the proanthocyanidin biosynthesis rather than the anthocyanin biosynthesis. Proanthocyanidin production could be an explanation for the identification of *DFR*, *ANS*, *LAR*, and *ANR* transcripts. However, it is important to emphasize that the transcript abundances of *DFR* and *ANS* are in any case extremely low (average TPM < 1), and inconsistent with high levels of anthocyanins. These low transcript abundances for DFR and ANS align well with previous reports about the absence of anthocyanin in betalain-pigmented species [8–10].

**Fig. 1:**
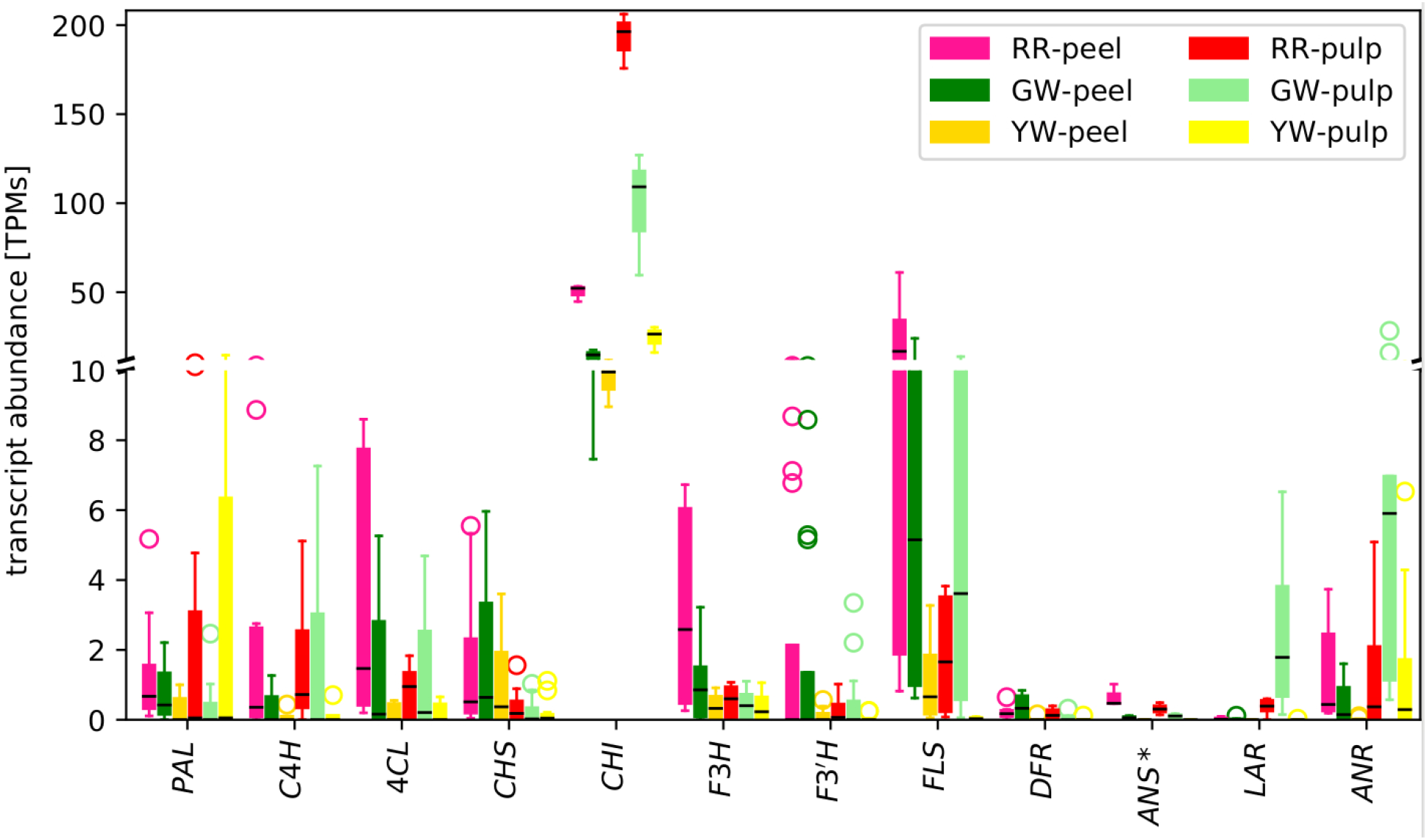
Transcript abundance of flavonoid biosynthesis genes in differently coloured pitaya cultivars. This transcript abundance analysis is based on our Trinity assembly of all RR datasets. No *bona fide* ANS was detected in the RR assembly, but the gene was represented with the closest homolog. RR=red peel/red pulp, GW=green peel/white pulp, YW= yellow peel/white pulp

## Conclusion

The evidence of anthocyanins lacks appropriate standards. Furthermore, the investigation of core anthocyanin biosynthesis genes via RNA-Seq does not provide insights into the accumulation of anthocyanins, because there is no clear difference in bulk anthocyanin content between differently pigmented pitaya varieties e.g. red vs. white. Consequently, there is little correlation between the transcription of anthocyanin synthesis genes, and proposed levels of anthocyanins. There is clear evidence of highly reduced DFR and ANS expression, which is not consistent with meaningful level so anthocyanins. Altogether, we suggest that the evidence of anthocyanins in pitaya remains weak.

## Methods

The applied methods are almost identical to our previous analysis of a very similar data set [7].

### Transcriptome assembly

RNAseq datasets of different cultivars were retrieved from the Sequence Read Archive via fastq-dump [11]. Trimming and adapter removal based on a set of all available Illumina adapters were performed via Trimmomatic v0.39 [12] using SLIDINGWINDOW:4:15 LEADING:5 TRAILING:5 MINLEN:50 TOPHRED33. A customized Python script was used to rename the surviving read pairs prior to the transcriptome assembly. Clean read pairs were subjected to Trinity v2.4.0 [13] for *de novo* transcriptome assembly using a k-mer size of 25. Short contigs below 200 bp were discarded. Previously described Python scripts [14] and BUSCO v3 [15] were applied for the calculation of assembly statistics for evaluation. Assembly quality was assessed based on continuity and completeness. Although assemblies were generated for all three species, the assembly generated on the basis of the data sets of *Hylocereus undatus* (SRR11603186-SRR11603191) was used for all down-stream analyses.

### Transcriptome annotation

Prediction of encoded peptides was performed using a previously described approach to identify and retain the longest predicted peptide per contig [14]. Genes involved in the flavonoid biosynthesis were identified via KIPEs [16] using the peptide mode. Phylogenetic trees with pitaya candidate sequences and previously characterized sequences [16] were constructed with FastTree v2 [17] (WAG+CAT model) based on alignments constructed via MAFFT v7 [18] and cleaned with pxclsq [19] to achieve a minimal occupancy of 0.1 for all alignment columns.

### Transcript abundance quantification

Quantification of transcript abundance was performed with kallisto v0.44.0 [20] using the RNAseq reads and our *Hylocereus undatus* transcriptome assembly [21]. Customized Python scripts were applied to summarize individual count tables and to compare expression values [7].

## ABBREVIATIONS

Not applicable

## ADDITIONAL FILES

Additional file 1: Phylogenetic trees of genes in the flavonoid biosynthesis. Identified candidate sequences are highlighted in red.

Additional file 2: Mapping table connecting the pitaya sequence names displayed in Additional file 1 to the original sequence IDs assigned during the assembly process.

## DECLARATIONS

### Ethics approval and consent to participate

Not applicable

### Consent for publication

Not applicable

### Availability of data and materials

The datasets generated and/or analysed during the current study are available via PUB: https://doi.org/10.4119/unibi/2956788. The Python scripts are available via github: https://github.com/bpucker/pitaya.

### Authors’ contributions

BP performed all analyses and prepared figures. BP and SFB wrote the manuscript. All authors reviewed the manuscript.

## Acknowledgements

We thank the Center for Biotechnology (CeBiTec) at Bielefeld University for providing an environment to perform the computational analyses. We thank Nathanael Walker-Hale for useful discussion.

