## Additional file 1 for "The evidence for anthocyanins in the betalain-pigmented genus Hylocereus is weak"

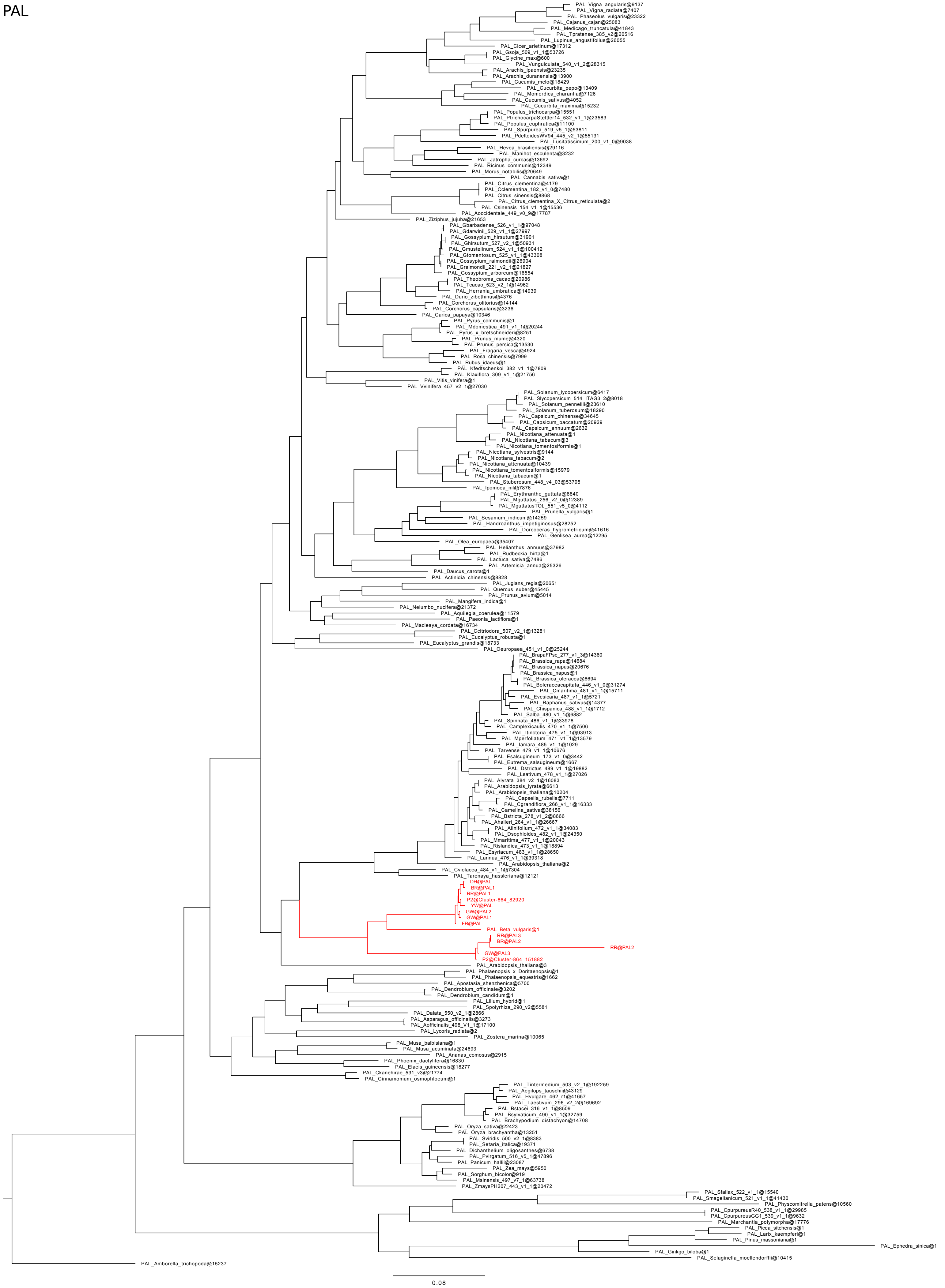
PAL
PAL_Vigna_angularis@9137
PAL_Vigna_radiata@7407
PAL_Phaseolus_vulgaris@23322
PAL_Cajanus_cajan@25083
PAL_Medicago_truncatula@41843
PAL_Tpratense_385_v2@20516
PAL_Lupinus_angustifolius@26055
PAL_Cicer_arietinum@17312
PAL_Gsoja_509_v1_1@53726
PAL_Glycine_max@600
PAL_Vunguiculata_540_v1_2@28315
PAL_Arachis_ipaensis@23235
PAL_Arachis_duranensis@13900
PAL_Cucumis_melo@18429
PAL_Cucurbita_pepo@13409
PAL_Momordica_charantia@7126
PAL_Cucumis_sativus@4052
PAL_Cucurbita_maxima@15232
PAL_Populus_trichocarpa@15551
PAL_PtrichocarpaStettler14_532_v1_1@23583
PAL_Populus_euphratica@11100
PAL_Spurpurea_519_v5_1@53811
PAL_PdeltoidesWV94_445_v2_1@55131
PAL_Lusitatissimum_200_v1_0@9038
PAL_Hevea_brasiliensis@29116
PAL_Manihot_esculenta@3232
PAL_Jatropha_curcas@13692
PAL_Ricinus_communis@12349
PAL_Morus_notabilis@20649
PAL_Cannabis_sativa@1
PAL_Citrus_clementina@4179
PAL_Cclementina_182_v1_0@7480
PAL_Citrus_sinensis@8868
PAL_Citrus_clementina_X_Citrus_reticulata@2
PAL_Csinensis_154_v1_1@15536
PAL_Aoccidentale_449_v0_9@17787
PAL_Ziziphus_jujuba@21653
PAL_Gbarbadense_526_v1_1@97048
PAL_Gdarwinii_529_v1_1@27997
PAL_Gossypium_hirsutum@31901
PAL_Ghirsutum_527_v2_1@50931
PAL_Gmustelinum_524_v1_1@100412
PAL_Gtomentosum_525_v1_1@43308
PAL_Gossypium_raimondii@26904
PAL_Graimondii_221_v2_1@21827
PAL_Gossypium_arboreum@16554
PAL_Theobroma_cacao@20986
PAL_Tcacao_523_v2_1@14962
PAL_Herrania_umbratica@14939
PAL_Durio_zibethinus@4376
PAL_Corchorus_olitorius@14144
PAL_Corchorus_capsularis@3236
PAL_Carica_papaya@10346
PAL_Pyrus_communis@1
PAL_Mdomestica_491_v1_1@20244
PAL_Pyrus_x_bretschneideri@8251
PAL_Prunus_mume@4320
PAL_Prunus_persica@13530
PAL_Fragaria_vesca@4924
PAL_Rosa_chinensis@7999
PAL_Rubus_idaeus@1
PAL_Kfedtschenkoi_382_v1_1@7809
PAL_Klaxiflora_309_v1_1@21756
PAL_Vitis_vinifera@1
PAL_Vvinifera_457_v2_1@27030
PAL_Solanum_lycopersicum@6417
PAL_Slycopersicum_514_ITAG3_2@8018
PAL_Solanum_pennellii@23610
PAL_Solanum_tuberosum@18290
PAL_Capsicum_chinense@34645
PAL_Capsicum_baccatum@20929
PAL_Capsicum_annuum@2632
PAL_Nicotiana_attenuata@1
PAL_Nicotiana_tabacum@3
PAL_Nicotiana_tomentosiformis@1
PAL_Nicotiana_sylvestris@9144
PAL_Nicotiana_tabacum@2
PAL_Nicotiana_attenuata@10439
PAL_Nicotiana_tomentosiformis@15979
PAL_Nicotiana_tabacum@1
PAL_Stuberosum_448_v4_03@53795
PAL_Ipomoea_nil@7876
PAL_Erythranthe_guttata@8840
PAL_Mguttatus_256_v2_0@12389
PAL_MguttatusTOL_551_v5_0@4112
PAL_Prunella_vulgaris@1
PAL_Sesamum_indicum@14259
PAL_Handroanthus_impetiginosus@28252
PAL_Dorcoceras_hygrometricum@41616
PAL_Genlisea_aurea@12295
PAL_Olea_europaea@35407
PAL_Helianthus_annuus@37982
PAL_Rudbeckia_hirta@1
PAL_Lactuca_sativa@7486
PAL_Artemisia_annua@25326
PAL_Daucus_carota@1
PAL_Actinidia_chinensis@8828
PAL_Juglans_regia@20651
PAL_Quercus_suber@45445
PAL_Prunus_avium@5014
PAL_Mangifera_indica@1
PAL_Nelumbo_nucifera@21372
PAL_Aquilegia_coerulea@11579
PAL_Paeonia_lactiflora@1
PAL_Macleaya_cordata@16734
PAL_Ccitriodora_507_v2_1@13281
PAL_Eucalyptus_robusta@1
PAL_Eucalyptus_grandis@18733
PAL_Oeuropaea_451_v1_0@25244
PAL_BrapaFPsc_277_v1_3@14360
PAL_Brassica_rapa@14684
PAL_Brassica_napus@20676
PAL_Brassica_napus@1
PAL_Brassica_oleracea@8694
PAL_Boleraceacapitata_446_v1_0@31274
PAL_Cmaritima_481_v1_1@15711
PAL_Evesicaria_487_v1_1@5721
PAL_Raphanus_sativus@14377
PAL_Chispanica_488_v1_1@1712
PAL_Salba_480_v1_1@6882
PAL_Spinnata_486_v1_1@33978
PAL_Camplexicaulis_470_v1_1@7506
PAL_Itinctoria_475_v1_1@93913
PAL_Mperfoliatum_471_v1_1@13579
PAL_Iamara_485_v1_1@1029
PAL_Tarvense_479_v1_1@10676
PAL_Esalsugineum_173_v1_0@3442
PAL_Eutrema_salsugineum@1667
PAL_Dstrictus_489_v1_1@19882
PAL_Lsativum_478_v1_1@27026
PAL_Alyrata_384_v2_1@16083
PAL_Arabidopsis_lyrata@6613
PAL_Arabidopsis_thaliana@10204
PAL_Capsella_rubella@7711
PAL_Cgrandiflora_266_v1_1@16333
PAL_Camelina_sativa@38156
PAL_Bstricta_278_v1_2@8666
PAL_Ahalleri_264_v1_1@26667
PAL_Alinifolium_472_v1_1@34083
PAL_Dsophioides_482_v1_1@24350
PAL_Mmaritima_477_v1_1@20043
PAL_Rislandica_473_v1_1@18894
PAL_Esyriacum_483_v1_1@28650
PAL_Lannua_476_v1_1@39318
PAL_Arabidopsis_thaliana@2
PAL_Cviolacea_484_v1_1@7304
PAL_Tarenaya_hassleriana@12121
DH@PAL
BR@PAL1
RR@PAL1
P2@Cluster-864_82920
YW@PAL
GW@PAL2
GW@PAL1
FR@PAL
PAL_Beta_vulgaris@1
RR@PAL3
BR@PAL2
RR@PAL2
GW@PAL3
P2@Cluster-864_151882
PAL_Arabidopsis_thaliana@3
PAL_Phalaenopsis_x_Doritaenopsis@1
PAL_Phalaenopsis_equestris@1662
PAL_Apostasia_shenzhenica@5700
PAL_Dendrobium_officinale@3202
PAL_Dendrobium_candidum@1
PAL_Lilium_hybrid@1
PAL_Spolyrhiza_290_v2@5581
PAL_Dalata_550_v2_1@2866
PAL_Asparagus_officinalis@3273
PAL_Aofficinalis_498_V1_1@17100
PAL_Lycoris_radiata@2
PAL_Zostera_marina@10065
PAL_Musa_balbisiana@1
PAL_Musa_acuminata@24693
PAL_Ananas_comosus@2915
PAL_Phoenix_dactylifera@16830
PAL_Elaeis_guineensis@18277
PAL_Ckanehirae_531_v3@21774
PAL_Cinnamomum_osmophloeum@1
PAL_Tintermedium_503_v2_1@192259
PAL_Aegilops_tauschii@43129
PAL_Hvulgare_462_r1@41657
PAL_Taestivum_296_v2_2@169692
PAL_Bstacei_316_v1_1@8509
PAL_Bsylvaticum_490_v1_1@32759
PAL_Brachypodium_distachyon@14708
PAL_Oryza_sativa@22423
PAL_Oryza_brachyantha@13251
PAL_Sviridis_500_v2_1@8383
PAL_Setaria_italica@19371
PAL_Dichanthelium_oligosanthes@6738
PAL_Pvirgatum_516_v5_1@47896
PAL_Panicum_hallii@23087
PAL_Zea_mays@5950
PAL_Sorghum_bicolor@919
PAL_Msinensis_497_v7_1@63738
PAL_ZmaysPH207_443_v1_1@20472
PAL_Sfallax_522_v1_1@15540
PAL_Smagellanicum_521_v1_1@41430
PAL_Physcomitrella_patens@10560
PAL_CpurpureusR40_538_v1_1@29985
PAL_CpurpureusGG1_539_v1_1@9632
PAL_Marchantia_polymorpha@17776
PAL_Picea_sitchensis@1
PAL_Larix_kaempferi@1
PAL_Pinus_massoniana@1
PAL_Ephedra_sinica@1
PAL_Ginkgo_biloba@1
PAL_Selaginella_moellendorffii@10415
PAL_Amborella_trichopoda@15237
0.08

C4H

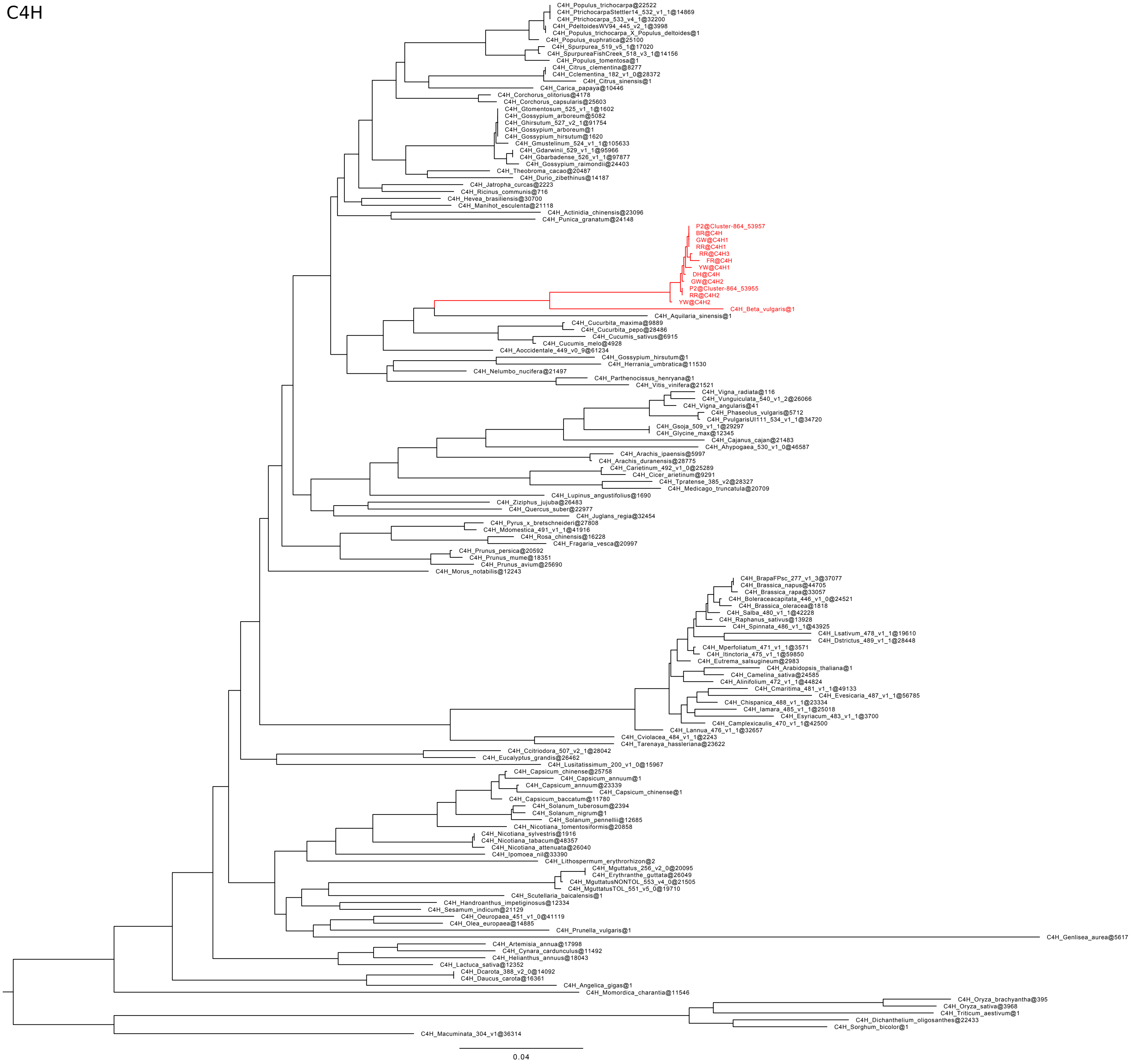

C4H_Populus_trichocarpa@22522
C4H_PtrichocarpaStettler14_532_v1_1@14869
C4H_Ptrichocarpa_533_v4_1@32200
C4H_PdeltoidesWV94_445_v2_1@3998
C4H_Populus_trichocarpa_X_Populus_deltoides@1
C4H_Populus_euphratica@25100
C4H_Spurpurea_519_v5_1@17020
C4H_SpurpureaFishCreek_518_v3_1@14156
C4H_Populus_tomentosa@1
C4H_Citrus_clementina@8277
C4H_Cclementina_182_v1_0@28372
C4H_Citrus_sinensis@1
C4H_Carica_papaya@10446
C4H_Corchorus_olitorius@4178
C4H_Corchorus_capsularis@25603
C4H_Gtomentosum_525_v1_1@1602
C4H_Gossypium_arboreum@5082
C4H_Ghirsutum_527_v2_1@91754
C4H_Gossypium_arboreum@1
C4H_Gossypium_hirsutum@1620
C4H_Gmustelinum_524_v1_1@105633
C4H_Gdarwinii_529_v1_1@95966
C4H_Gbarbadense_526_v1_1@97877
C4H_Gossypium_raimondii@24403
C4H_Theobroma_cacao@20487
C4H_Durio_zibethinus@14187
C4H_Jatropha_curcas@2223
C4H_Ricinus_communis@716
C4H_Hevea_brasiliensis@30700
C4H_Manihot_esculenta@21118
C4H_Actinidia_chinensis@23096
C4H_Punica_granatum@24148
P2@Cluster-864_53957
BR@C4H
GW@C4H1
RR@C4H1
RR@C4H3
FR@C4H
YW@C4H1
DH@C4H
GW@C4H2
P2@Cluster-864_53955
RR@C4H2
YW@C4H2
C4H_Beta_vulgaris@1
C4H_Aquilaria_sinensis@1
C4H_Cucurbita_maxima@9889
C4H_Cucurbita_pepo@28486
C4H_Cucumis_sativus@6915
C4H_Cucumis_melo@4928
C4H_Aoccidentale_449_v0_9@61234
C4H_Gossypium_hirsutum@1
C4H_Herrania_umbratica@11530
C4H_Nelumbo_nucifera@21497
C4H_Parthenocissus_henryana@1
C4H_Vitis_vinifera@21521
C4H_Vigna_radiata@116
C4H_Vunguiculata_540_v1_2@26066
C4H_Vigna_angularis@41
C4H_Phaseolus_vulgaris@5712
C4H_PvulgarisUI111_534_v1_1@34720
C4H_Gsoja_509_v1_1@29297
C4H_Glycine_max@12345
C4H_Cajanus_cajan@21483
C4H_Ahypogaea_530_v1_0@46587
C4H_Arachis_ipaensis@5997
C4H_Arachis_duranensis@28775
C4H_Carietinum_492_v1_0@25289
C4H_Cicer_arietinum@9291
C4H_Tpratense_385_v2@28327
C4H_Medicago_truncatula@20709
C4H_Lupinus_angustifolius@1690
C4H_Ziziphus_jujuba@26483
C4H_Quercus_suber@22977
C4H_Juglans_regia@32454
C4H_Pyrus_x_bretschneideri@27808
C4H_Mdomestica_491_v1_1@41916
C4H_Rosa_chinensis@16228
C4H_Fragaria_vesca@20997
C4H_Prunus_persica@20592
C4H_Prunus_mume@18351
C4H_Prunus_avium@25690
C4H_Morus_notabilis@12243
C4H_BrapaFPsc_277_v1_3@37077
C4H_Brassica_napus@44705
C4H_Brassica_rapa@33057
C4H_Boleraceacapitata_446_v1_0@24521
C4H_Brassica_oleracea@1818
C4H_Salba_480_v1_1@42228
C4H_Raphanus_sativus@13928
C4H_Spinnata_486_v1_1@43925
C4H_Lsativum_478_v1_1@19610
C4H_Dstrictus_489_v1_1@28448
C4H_Mperfoliatum_471_v1_1@3571
C4H_Itinctoria_475_v1_1@59850
C4H_Eutrema_salsugineum@2983
C4H_Arabidopsis_thaliana@1
C4H_Camelina_sativa@24585
C4H_Alinifolium_472_v1_1@44824
C4H_Cmaritima_481_v1_1@49133
C4H_Evesicaria_487_v1_1@56785
C4H_Chispanica_488_v1_1@23334
C4H_Iamara_485_v1_1@25018
C4H_Esyriacum_483_v1_1@3700
C4H_Camplexicaulis_470_v1_1@42500
C4H_Lannua_476_v1_1@32657
C4H_Cviolacea_484_v1_1@2243
C4H_Tarenaya_hassleriana@23622
C4H_Ccitriodora_507_v2_1@28042
C4H_Eucalyptus_grandis@26462
C4H_Lusitatissimum_200_v1_0@15967
C4H_Capsicum_chinense@25758
C4H_Capsicum_annuum@1
C4H_Capsicum_annuum@23339
C4H_Capsicum_chinense@1
C4H_Capsicum_baccatum@11780
C4H_Solanum_tuberosum@2394
C4H_Solanum_nigrum@1
C4H_Solanum_pennellii@12685
C4H_Nicotiana_tomentosiformis@20858
C4H_Nicotiana_sylvestris@1916
C4H_Nicotiana_tabacum@48357
C4H_Nicotiana_attenuata@26040
C4H_Ipomoea_nil@33390
C4H_Lithospermum_erythrorhizon@2
C4H_Mguttatus_256_v2_0@20095
C4H_Erythranthe_guttata@26049
C4H_MguttatusNONTOL_553_v4_0@21505
C4H_MguttatusTOL_551_v5_0@19710
C4H_Scutellaria_baicalensis@1
C4H_Handroanthus_impetiginosus@12334
C4H_Sesamum_indicum@21129
C4H_Oeuropaea_451_v1_0@41119
C4H_Olea_europaea@14885
C4H_Prunella_vulgaris@1
C4H_Genlisea_aurea@5617
C4H_Artemisia_annua@17998
C4H_Cynara_cardunculus@11492
C4H_Helianthus_annuus@18043
C4H_Lactuca_sativa@12352
C4H_Dcarota_388_v2_0@14092
C4H_Daucus_carota@16361
C4H_Angelica_gigas@1
C4H_Momordica_charantia@11546
C4H_Oryza_brachyantha@395
C4H_Oryza_sativa@3968
C4H_Triticum_aestivum@1
C4H_Dichanthelium_oligosanthes@22433
C4H_Sorghum_bicolor@1
C4H_Macuminata_304_v1@36314
0.04

4CL

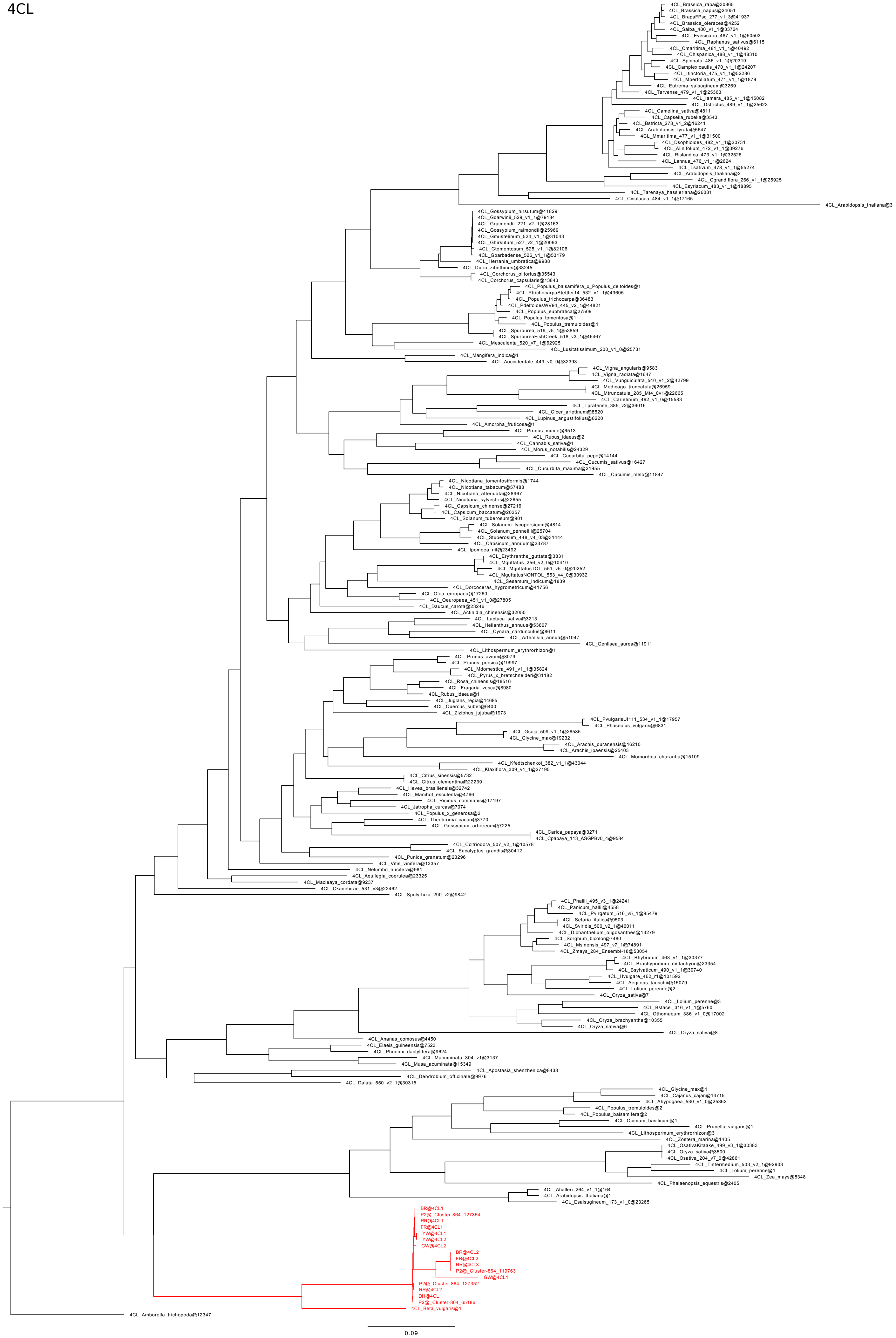

4CL_Brassica_rapa@30865
4CL_Brassica_napus@24051
4CL_BrapaFPsc_277_v1_3@41937
4CL_Brassica_oleracea@4252
4CL_Salba_480_v1_1@33724
4CL_Evesicaria_487_v1_1@50503
4CL_Raphanus_sativus@6115
4CL_Cmaritima_481_v1_1@40492
4CL_Chispanica_488_v1_1@48310
4CL_Spinnata_486_v1_1@20319
4CL_Camplexicaulis_470_v1_1@24207
4CL_Itinctoria_475_v1_1@52286
4CL_Mperfoliatum_471_v1_1@1879
4CL_Eutrema_salsugineum@3269
4CL_Tarvense_479_v1_1@25363
4CL_Iamara_485_v1_1@15082
4CL_Dstrictus_489_v1_1@25623
4CL_Camelina_sativa@4811
4CL_Capsella_rubella@3543
4CL_Bstricta_278_v1_2@16241
4CL_Arabidopsis_lyrata@5647
4CL_Mmaritima_477_v1_1@31500
4CL_Dsophioides_482_v1_1@20731
4CL_Alinifolium_472_v1_1@39276
4CL_Rislandica_473_v1_1@32526
4CL_Lannua_476_v1_1@2624
4CL_Lsativum_478_v1_1@55274
4CL_Arabidopsis_thaliana@2
4CL_Cgrandiflora_266_v1_1@25925
4CL_Esyriacum_483_v1_1@18895
4CL_Tarenaya_hassleriana@26081
4CL_Cviolacea_484_v1_1@17165
4CL_Arabidopsis_thaliana@3
4CL_Gossypium_hirsutum@41829
4CL_Gdarwinii_529_v1_1@79184
4CL_Graimondii_221_v2_1@28163
4CL_Gossypium_raimondii@25969
4CL_Gmustelinum_524_v1_1@31043
4CL_Ghirsutum_527_v2_1@20093
4CL_Gtomentosum_525_v1_1@82106
4CL_Gbarbadense_526_v1_1@53179
4CL_Herrania_umbratica@9988
4CL_Durio_zibethinus@33245
4CL_Corchorus_olitorius@35543
4CL_Corchorus_capsularis@13843
4CL_Populus_balsamifera_x_Populus_deltoides@1
4CL_PtrichocarpaStettler14_532_v1_1@49605
4CL_Populus_trichocarpa@36483
4CL_PdeltoidesWV94_445_v2_1@44821
4CL_Populus_euphratica@27509
4CL_Populus_tomentosa@1
4CL_Populus_tremuloides@1
4CL_Spurpurea_519_v5_1@53859
4CL_SpurpureaFishCreek_518_v3_1@46467
4CL_Mesculenta_520_v7_1@62925
4CL_Lusitatissimum_200_v1_0@25731
4CL_Mangifera_indica@1
4CL_Aoccidentale_449_v0_9@32393
4CL_Vigna_angularis@9583
4CL_Vigna_radiata@1647
4CL_Vunguiculata_540_v1_2@42799
4CL_Medicago_truncatula@26959
4CL_Mtruncatula_285_Mt4_0v1@22665
4CL_Carietinum_492_v1_0@15583
4CL_Tpratense_385_v2@36016
4CL_Cicer_arietinum@8520
4CL_Lupinus_angustifolius@6220
4CL_Amorpha_fruticosa@1
4CL_Prunus_mume@6513
4CL_Rubus_idaeus@2
4CL_Cannabis_sativa@1
4CL_Morus_notabilis@24329
4CL_Cucurbita_pepo@14144
4CL_Cucumis_sativus@16427
4CL_Cucurbita_maxima@21955
4CL_Cucumis_melo@11847
4CL_Nicotiana_tomentosiformis@1744
4CL_Nicotiana_tabacum@57488
4CL_Nicotiana_attenuata@28967
4CL_Nicotiana_sylvestris@22655
4CL_Capsicum_chinense@27216
4CL_Capsicum_baccatum@20257
4CL_Solanum_tuberosum@901
4CL_Solanum_lycopersicum@4814
4CL_Solanum_pennellii@25704
4CL_Stuberosum_448_v4_03@31444
4CL_Capsicum_annuum@23787
4CL_Ipomoea_nil@23492
4CL_Erythranthe_guttata@3831
4CL_Mguttatus_256_v2_0@10410
4CL_MguttatusTOL_551_v5_0@20252
4CL_MguttatusNONTOL_553_v4_0@30932
4CL_Sesamum_indicum@1839
4CL_Dorcoceras_hygrometricum@41756
4CL_Olea_europaea@17260
4CL_Oeuropaea_451_v1_0@27805
4CL_Daucus_carota@23246
4CL_Actinidia_chinensis@32050
4CL_Lactuca_sativa@3213
4CL_Helianthus_annuus@53807
4CL_Cynara_cardunculus@8611
4CL_Artemisia_annua@51047
4CL_Genlisea_aurea@11911
4CL_Lithospermum_erythrorhizon@1
4CL_Prunus_avium@8079
4CL_Prunus_persica@19997
4CL_Mdomestica_491_v1_1@35824
4CL_Pyrus_x_bretschneideri@31182
4CL_Rosa_chinensis@18516
4CL_Fragaria_vesca@8980
4CL_Rubus_idaeus@1
4CL_Juglans_regia@14685
4CL_Quercus_suber@6400
4CL_Ziziphus_jujuba@1973
4CL_PvulgarisUI111_534_v1_1@17957
4CL_Phaseolus_vulgaris@6831
4CL_Gsoja_509_v1_1@28585
4CL_Glycine_max@19232
4CL_Arachis_duranensis@16210
4CL_Arachis_ipaensis@25403
4CL_Momordica_charantia@15109
4CL_Kfedtschenkoi_382_v1_1@43044
4CL_Klaxiflora_309_v1_1@27195
4CL_Citrus_sinensis@5732
4CL_Citrus_clementina@22239
4CL_Hevea_brasiliensis@32742
4CL_Manihot_esculenta@4766
4CL_Ricinus_communis@17197
4CL_Jatropha_curcas@7074
4CL_Populus_x_generosa@2
4CL_Theobroma_cacao@3770
4CL_Gossypium_arboreum@7225
4CL_Carica_papaya@3271
4CL_Cpapaya_113_ASGPBv0_4@9584
4CL_Ccitriodora_507_v2_1@10578
4CL_Eucalyptus_grandis@30412
4CL_Punica_granatum@23296
4CL_Vitis_vinifera@13357
4CL_Nelumbo_nucifera@981
4CL_Aquilegia_coerulea@23325
4CL_Macleaya_cordata@9237
4CL_Ckanehirae_531_v3@22462
4CL_Spolyrhiza_290_v2@9842
4CL_Phallii_495_v3_1@24241
4CL_Panicum_hallii@4558
4CL_Pvirgatum_516_v5_1@95479
4CL_Setaria_italica@9503
4CL_Sviridis_500_v2_1@46011
4CL_Dichanthelium_oligosanthes@13279
4CL_Sorghum_bicolor@7480
4CL_Msinensis_497_v7_1@74891
4CL_Zmays_284_Ensembl-18@53054
4CL_Bhybridum_463_v1_1@30377
4CL_Brachypodium_distachyon@23354
4CL_Bsylvaticum_490_v1_1@39740
4CL_Hvulgare_462_r1@101592
4CL_Aegilops_tauschii@15079
4CL_Lolium_perenne@2
4CL_Oryza_sativa@7
4CL_Lolium_perenne@3
4CL_Bstacei_316_v1_1@5760
4CL_Othomaeum_386_v1_0@17002
4CL_Oryza_brachyantha@10355
4CL_Oryza_sativa@6
4CL_Oryza_sativa@8
4CL_Ananas_comosus@4450
4CL_Elaeis_guineensis@7523
4CL_Phoenix_dactylifera@9624
4CL_Macuminata_304_v1@3137
4CL_Musa_acuminata@15349
4CL_Apostasia_shenzhenica@8438
4CL_Dendrobium_officinale@9976
4CL_Dalata_550_v2_1@30315
4CL_Glycine_max@1
4CL_Cajanus_cajan@14715
4CL_Ahypogaea_530_v1_0@25362
4CL_Populus_tremuloides@2
4CL_Populus_balsamifera@2
4CL_Ocimum_basilicum@1
4CL_Prunella_vulgaris@1
4CL_Lithospermum_erythrorhizon@3
4CL_Zostera_marina@1405
4CL_OsativaKitaake_499_v3_1@30383
4CL_Oryza_sativa@3500
4CL_Osativa_204_v7_0@42861
4CL_Tintermedium_503_v2_1@92903
4CL_Lolium_perenne@1
4CL_Zea_mays@8348
4CL_Phalaenopsis_equestris@2405
4CL_Ahalleri_264_v1_1@164
4CL_Arabidopsis_thaliana@1
4CL_Esalsugineum_173_v1_0@23265
BR@4CL1
P2@_Cluster-864_127354
RR@4CL1
FR@4CL1
YW@4CL1
YW@4CL2
GW@4CL2
BR@4CL2
FR@4CL2
RR@4CL3
P2@_Cluster-864_119763
GW@4CL1
P2@_Cluster-864_127352
RR@4CL2
DH@4CL
P2@_Cluster-864_65186
4CL_Beta_vulgaris@1
4CL_Amborella_trichopoda@12347
0.09

### CHS

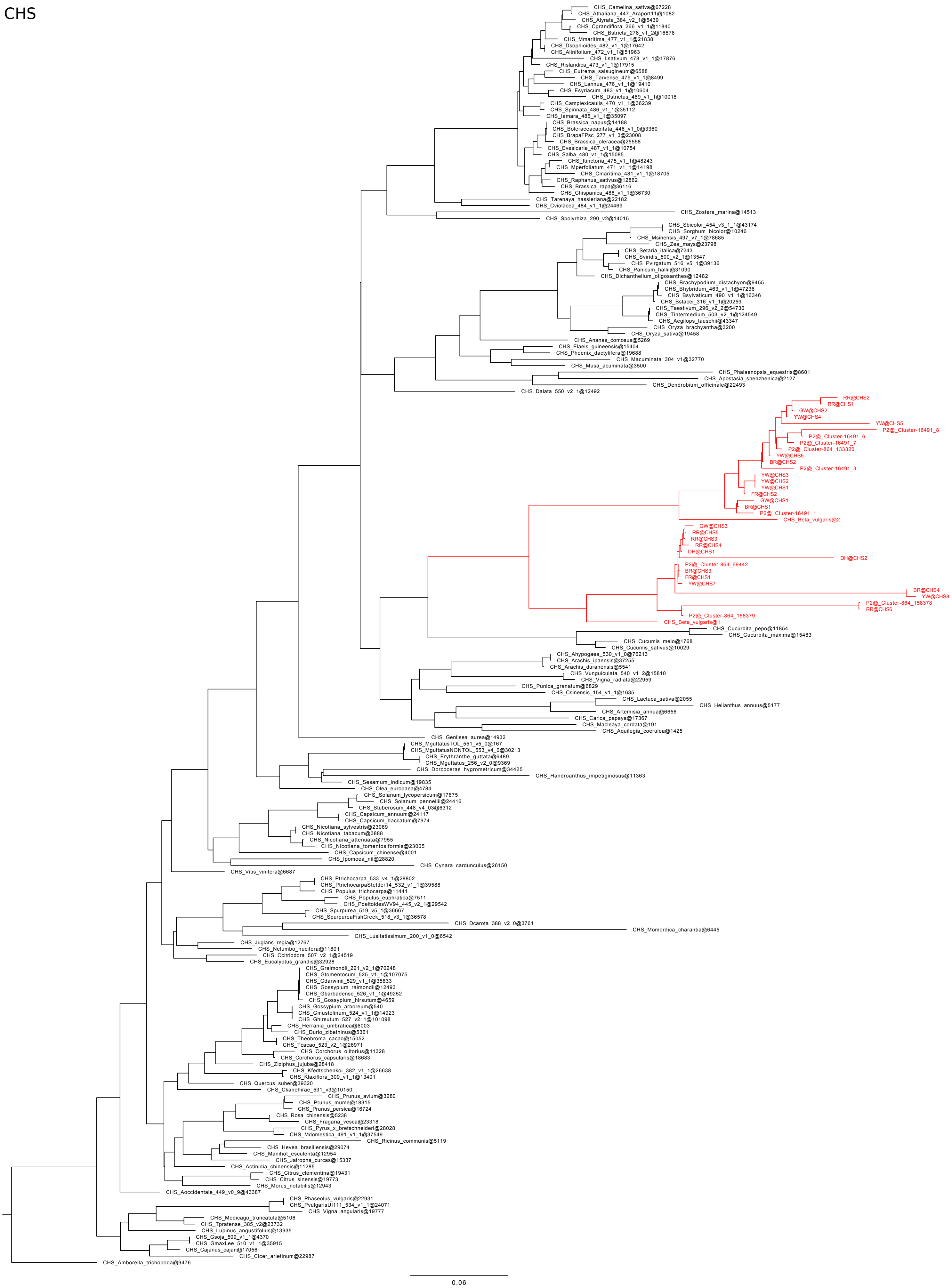
CHS_Camelina_sativa@67228
CHS_Athaliana_447_Araport11@1082
CHS_Alyrata_384_v2_1@5439
CHS_Cgrandiflora_266_v1_1@11840
CHS_Bstricta_278_v1_2@16878
CHS_Mmaritima_477_v1_1@21838
CHS_Dsophioides_482_v1_1@17642
CHS_Alinifolium_472_v1_1@51963
CHS_Lsativum_478_v1_1@17876
CHS_Rislandica_473_v1_1@17915
CHS_Eutrema_salsugineum@6588
CHS_Tarvense_479_v1_1@8499
CHS_Lannua_476_v1_1@19410
CHS_Esyriacum_483_v1_1@10604
CHS_Dstrictus_489_v1_1@10018
CHS_Camplexicaulis_470_v1_1@36239
CHS_Spinnata_486_v1_1@35112
CHS_Iamara_485_v1_1@35097
CHS_Brassica_napus@14188
CHS_Boleraceacapitata_446_v1_0@3360
CHS_BrapaFPsc_277_v1_3@23008
CHS_Brassica_oleracea@25558
CHS_Evesicaria_487_v1_1@10754
CHS_Salba_480_v1_1@15085
CHS_Itinctoria_475_v1_1@48243
CHS_Mperfoliatum_471_v1_1@14198
CHS_Cmaritima_481_v1_1@18705
CHS_Raphanus_sativus@12862
CHS_Brassica_rapa@36116
CHS_Chispanica_488_v1_1@36730
CHS_Tarenaya_hassleriana@22182
CHS_Cviolacea_484_v1_1@24469
CHS_Zostera_marina@14513
CHS_Spolyrhiza_290_v2@14015
CHS_Sbicolor_454_v3_1_1@43174
CHS_Sorghum_bicolor@10246
CHS_Msinensis_497_v7_1@78685
CHS_Zea_mays@23798
CHS_Setaria_italica@7243
CHS_Sviridis_500_v2_1@13547
CHS_Pvirgatum_516_v5_1@39136
CHS_Panicum_hallii@31090
CHS_Dichanthelium_oligosanthes@12482
CHS_Brachypodium_distachyon@9455
CHS_Bhybridum_463_v1_1@47236
CHS_Bsylvaticum_490_v1_1@16346
CHS_Bstacei_316_v1_1@20259
CHS_Taestivum_296_v2_2@54730
CHS_Tintermedium_503_v2_1@124549
CHS_Aegilops_tauschii@43347
CHS_Oryza_brachyantha@3200
CHS_Oryza_sativa@19458
CHS_Ananas_comosus@5269
CHS_Elaeis_guineensis@15404
CHS_Phoenix_dactylifera@19688
CHS_Macuminata_304_v1@32770
CHS_Musa_acuminata@3500
CHS_Phalaenopsis_equestris@8601
CHS_Apostasia_shenzhenica@2127
CHS_Dendrobium_officinale@22493
CHS_Dalata_550_v2_1@12492
RR@CHS2
RR@CHS1
GW@CHS2
YW@CHS4
YW@CHS5
P2@_Cluster-16491_8
P2@_Cluster-16491_6
P2@_Cluster-16491_7
P2@_Cluster-864_133320
YW@CHS6
BR@CHS2
P2@_Cluster-16491_3
YW@CHS3
YW@CHS2
YW@CHS1
FR@CHS2
GW@CHS1
BR@CHS1
P2@_Cluster-16491_1
CHS_Beta_vulgaris@2
GW@CHS3
RR@CHS5
RR@CHS3
RR@CHS4
DH@CHS1
DH@CHS2
P2@_Cluster-864_69442
BR@CHS3
FR@CHS1
YW@CHS7
BR@CHS4
YW@CHS8
P2@_Cluster-864_158378
RR@CHS6
P2@_Cluster-864_158379
CHS_Beta_vulgaris@1
CHS_Cucurbita_pepo@11854
CHS_Cucurbita_maxima@15483
CHS_Cucumis_melo@1768
CHS_Cucumis_sativus@10029
CHS_Ahypogaea_530_v1_0@76213
CHS_Arachis_ipaensis@37255
CHS_Arachis_duranensis@5541
CHS_Vunguiculata_540_v1_2@15810
CHS_Vigna_radiata@22959
CHS_Punica_granatum@6829
CHS_Csinensis_154_v1_1@1635
CHS_Lactuca_sativa@2055
CHS_Helianthus_annuus@5177
CHS_Artemisia_annua@6656
CHS_Carica_papaya@17367
CHS_Macleaya_cordata@191
CHS_Aquilegia_coerulea@1425
CHS_Genlisea_aurea@14932
CHS_MguttatusTOL_551_v5_0@167
CHS_MguttatusNONTOL_553_v4_0@30213
CHS_Erythranthe_guttata@6489
CHS_Mguttatus_256_v2_0@9369
CHS_Dorcoceras_hygrometricum@34425
CHS_Handroanthus_impetiginosus@11363
CHS_Sesamum_indicum@19835
CHS_Olea_europaea@4784
CHS_Solanum_lycopersicum@17675
CHS_Solanum_pennellii@24416
CHS_Stuberosum_448_v4_03@6312
CHS_Capsicum_annuum@24117
CHS_Capsicum_baccatum@7974
CHS_Nicotiana_sylvestris@23069
CHS_Nicotiana_tabacum@3888
CHS_Nicotiana_attenuata@7955
CHS_Nicotiana_tomentosiformis@23005
CHS_Capsicum_chinense@4001
CHS_Ipomoea_nil@28820
CHS_Cynara_cardunculus@26150
CHS_Vitis_vinifera@6687
CHS_Ptrichocarpa_533_v4_1@28802
CHS_PtrichocarpaStettler14_532_v1_1@39588
CHS_Populus_trichocarpa@11441
CHS_Populus_euphratica@7511
CHS_PdeltoidesWV94_445_v2_1@29542
CHS_Spurpurea_519_v5_1@36667
CHS_SpurpureaFishCreek_518_v3_1@36578
CHS_Dcarota_388_v2_0@3761
CHS_Momordica_charantia@6445
CHS_Lusitatissimum_200_v1_0@6542
CHS_Juglans_regia@12767
CHS_Nelumbo_nucifera@11801
CHS_Ccitriodora_507_v2_1@24519
CHS_Eucalyptus_grandis@32928
CHS_Graimondii_221_v2_1@70248
CHS_Gtomentosum_525_v1_1@107075
CHS_Gdarwinii_529_v1_1@35833
CHS_Gossypium_raimondii@12493
CHS_Gbarbadense_526_v1_1@49252
CHS_Gossypium_hirsutum@4659
CHS_Gossypium_arboreum@540
CHS_Gmustelinum_524_v1_1@14923
CHS_Ghirsutum_527_v2_1@101098
CHS_Herrania_umbratica@6003
CHS_Durio_zibethinus@5361
CHS_Theobroma_cacao@15052
CHS_Tcacao_523_v2_1@26971
CHS_Corchorus_olitorius@11328
CHS_Corchorus_capsularis@18683
CHS_Ziziphus_jujuba@28418
CHS_Kfedtschenkoi_382_v1_1@26638
CHS_Klaxiflora_309_v1_1@13401
CHS_Quercus_suber@39320
CHS_Ckanehirae_531_v3@10150
CHS_Prunus_avium@3280
CHS_Prunus_mume@18315
CHS_Prunus_persica@16724
CHS_Rosa_chinensis@5238
CHS_Fragaria_vesca@23318
CHS_Pyrus_x_bretschneideri@28028
CHS_Mdomestica_491_v1_1@37549
CHS_Ricinus_communis@5119
CHS_Hevea_brasiliensis@29074
CHS_Manihot_esculenta@12954
CHS_Jatropha_curcas@15337
CHS_Actinidia_chinensis@11285
CHS_Citrus_clementina@19431
CHS_Citrus_sinensis@19773
CHS_Morus_notabilis@12943
CHS_Aoccidentale_449_v0_9@43387
CHS_Phaseolus_vulgaris@22931
CHS_PvulgarisUI111_534_v1_1@24071
CHS_Vigna_angularis@19777
CHS_Medicago_truncatula@5106
CHS_Tpratense_385_v2@23732
CHS_Lupinus_angustifolius@13935
CHS_Gsoja_509_v1_1@4370
CHS_GmaxLee_510_v1_1@35915
CHS_Cajanus_cajan@17056
CHS_Cicer_arietinum@22987
CHS_Amborella_trichopoda@9476
0.06

CHI

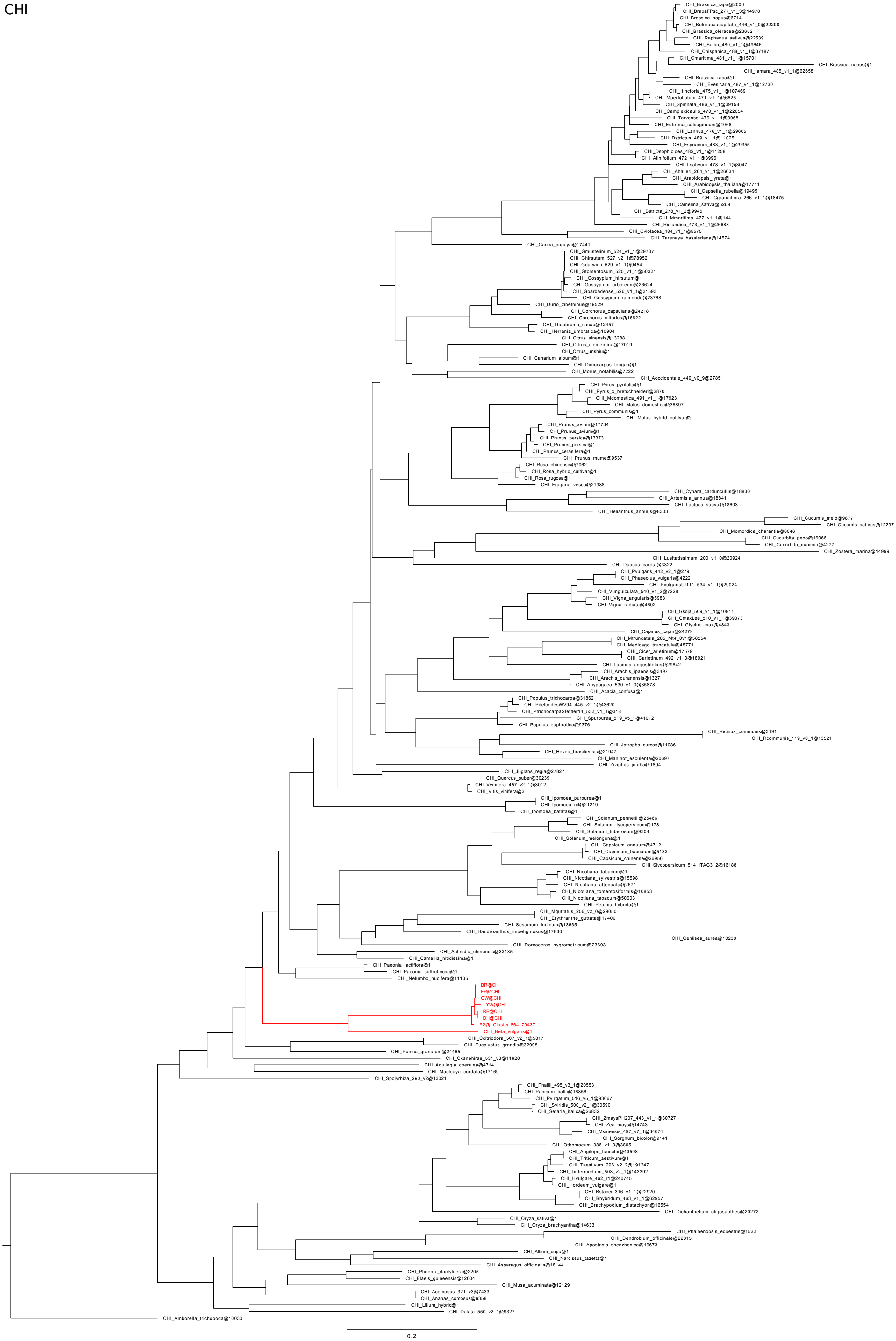

CHI_Brassica_rapa@2006
CHI_BrapaFPsc_277_v1_3@14978
CHI_Brassica_napus@67141
CHI_Boleraceacapitata_446_v1_0@22298
CHI_Brassica_oleracea@23652
CHI_Raphanus_sativus@22539
CHI_Salba_480_v1_1@49646
CHI_Chispanica_488_v1_1@37187
CHI_Cmaritima_481_v1_1@15701
CHI_Brassica_napus@1
CHI_Iamara_485_v1_1@62658
CHI_Brassica_rapa@1
CHI_Evesicaria_487_v1_1@12730
CHI_Itinctoria_475_v1_1@107469
CHI_Mperfoliatum_471_v1_1@6625
CHI_Spinnata_486_v1_1@39158
CHI_Camplexicaulis_470_v1_1@22054
CHI_Tarvense_479_v1_1@3068
CHI_Eutrema_salsugineum@4068
CHI_Lannua_476_v1_1@29605
CHI_Dstrictus_489_v1_1@11025
CHI_Esyriacum_483_v1_1@29355
CHI_Dsophioides_482_v1_1@11258
CHI_Alinifolium_472_v1_1@39961
CHI_Lsativum_478_v1_1@3047
CHI_Ahalleri_264_v1_1@26634
CHI_Arabidopsis_lyrata@1
CHI_Arabidopsis_thaliana@17711
CHI_Capsella_rubella@19495
CHI_Cgrandiflora_266_v1_1@18475
CHI_Camelina_sativa@5269
CHI_Bstricta_278_v1_2@9945
CHI_Mmaritima_477_v1_1@144
CHI_Rislandica_473_v1_1@26688
CHI_Cviolacea_484_v1_1@5575
CHI_Tarenaya_hassleriana@14574
CHI_Carica_papaya@17441
CHI_Gmustelinum_524_v1_1@29707
CHI_Ghirsutum_527_v2_1@78952
CHI_Gdarwinii_529_v1_1@9454
CHI_Gtomentosum_525_v1_1@50321
CHI_Gossypium_hirsutum@1
CHI_Gossypium_arboreum@26624
CHI_Gbarbadense_526_v1_1@31593
CHI_Gossypium_raimondii@23768
CHI_Durio_zibethinus@19529
CHI_Corchorus_capsularis@24218
CHI_Corchorus_olitorius@16822
CHI_Theobroma_cacao@12457
CHI_Herrania_umbratica@10904
CHI_Citrus_sinensis@13288
CHI_Citrus_clementina@17019
CHI_Citrus_unshiu@1
CHI_Canarium_album@1
CHI_Dimocarpus_longan@1
CHI_Morus_notabilis@7222
CHI_Aoccidentale_449_v0_9@27851
CHI_Pyrus_pyrifolia@1
CHI_Pyrus_x_bretschneideri@2870
CHI_Mdomestica_491_v1_1@17923
CHI_Malus_domestica@36897
CHI_Pyrus_communis@1
CHI_Malus_hybrid_cultivar@1
CHI_Prunus_avium@17734
CHI_Prunus_avium@1
CHI_Prunus_persica@13373
CHI_Prunus_persica@1
CHI_Prunus_cerasifera@1
CHI_Prunus_mume@9537
CHI_Rosa_chinensis@7062
CHI_Rosa_hybrid_cultivar@1
CHI_Rosa_rugosa@1
CHI_Fragaria_vesca@21988
CHI_Cynara_cardunculus@18830
CHI_Artemisia_annua@18841
CHI_Lactuca_sativa@18603
CHI_Helianthus_annuus@8303
CHI_Cucumis_melo@9877
CHI_Cucumis_sativus@12297
CHI_Momordica_charantia@6646
CHI_Cucurbita_pepo@16066
CHI_Cucurbita_maxima@4277
CHI_Zostera_marina@14999
CHI_Lusitatissimum_200_v1_0@20924
CHI_Daucus_carota@3322
CHI_Pvulgaris_442_v2_1@279
CHI_Phaseolus_vulgaris@4222
CHI_PvulgarisUI111_534_v1_1@29024
CHI_Vunguiculata_540_v1_2@7228
CHI_Vigna_angularis@5988
CHI_Vigna_radiata@4602
CHI_Gsoja_509_v1_1@10911
CHI_GmaxLee_510_v1_1@39373
CHI_Glycine_max@4843
CHI_Cajanus_cajan@24279
CHI_Mtruncatula_285_Mt4_0v1@58254
CHI_Medicago_truncatula@48771
CHI_Cicer_arietinum@17579
CHI_Carietinum_492_v1_0@18921
CHI_Lupinus_angustifolius@29842
CHI_Arachis_ipaensis@3497
CHI_Arachis_duranensis@1327
CHI_Ahypogaea_530_v1_0@35878
CHI_Acacia_confusa@1
CHI_Populus_trichocarpa@31862
CHI_PdeltoidesWV94_445_v2_1@43620
CHI_PtrichocarpaStettler14_532_v1_1@318
CHI_Spurpurea_519_v5_1@41012
CHI_Populus_euphratica@9376
CHI_Ricinus_communis@3191
CHI_Rcommunis_119_v0_1@13521
CHI_Jatropha_curcas@11086
CHI_Hevea_brasiliensis@21947
CHI_Manihot_esculenta@20697
CHI_Ziziphus_jujuba@1894
CHI_Juglans_regia@27827
CHI_Quercus_suber@30239
CHI_Vvinifera_457_v2_1@3012
CHI_Vitis_vinifera@2
CHI_Ipomoea_purpurea@1
CHI_Ipomoea_nil@21219
CHI_Ipomoea_batatas@1
CHI_Solanum_pennellii@25466
CHI_Solanum_lycopersicum@178
CHI_Solanum_tuberosum@9304
CHI_Solanum_melongena@1
CHI_Capsicum_annuum@4712
CHI_Capsicum_baccatum@5182
CHI_Capsicum_chinense@26956
CHI_Slycopersicum_514_ITAG3_2@16188
CHI_Nicotiana_tabacum@1
CHI_Nicotiana_sylvestris@15598
CHI_Nicotiana_attenuata@2671
CHI_Nicotiana_tomentosiformis@10853
CHI_Nicotiana_tabacum@50003
CHI_Petunia_hybrida@1
CHI_Mguttatus_256_v2_0@29050
CHI_Erythranthe_guttata@17400
CHI_Sesamum_indicum@13635
CHI_Handroanthus_impetiginosus@17830
CHI_Genlisea_aurea@10238
CHI_Dorcoceras_hygrometricum@23693
CHI_Actinidia_chinensis@32185
CHI_Camellia_nitidissima@1
CHI_Paeonia_lactiflora@1
CHI_Paeonia_suffruticosa@1
CHI_Nelumbo_nucifera@11135
BR@CHI
FR@CHI
GW@CHI
YW@CHI
RR@CHI
DH@CHI
P2@_Cluster-864_79437
CHI_Beta_vulgaris@1
CHI_Ccitriodora_507_v2_1@5817
CHI_Eucalyptus_grandis@32998
CHI_Punica_granatum@24465
CHI_Ckanehirae_531_v3@11920
CHI_Aquilegia_coerulea@4714
CHI_Macleaya_cordata@17169
CHI_Spolyrhiza_290_v2@13021
CHI_Phallii_495_v3_1@20553
CHI_Panicum_hallii@16856
CHI_Pvirgatum_516_v5_1@93667
CHI_Sviridis_500_v2_1@30590
CHI_Setaria_italica@26832
CHI_ZmaysPH207_443_v1_1@30727
CHI_Zea_mays@14743
CHI_Msinensis_497_v7_1@34674
CHI_Sorghum_bicolor@9141
CHI_Othomaeum_386_v1_0@3805
CHI_Aegilops_tauschii@43598
CHI_Triticum_aestivum@1
CHI_Taestivum_296_v2_2@191247
CHI_Tintermedium_503_v2_1@143392
CHI_Hvulgare_462_r1@240745
CHI_Hordeum_vulgare@1
CHI_Bstacei_316_v1_1@22920
CHI_Bhybridum_463_v1_1@62957
CHI_Brachypodium_distachyon@16554
CHI_Dichanthelium_oligosanthes@20272
CHI_Oryza_sativa@1
CHI_Oryza_brachyantha@14633
CHI_Phalaenopsis_equestris@1522
CHI_Dendrobium_officinale@22815
CHI_Apostasia_shenzhenica@19673
CHI_Allium_cepa@1
CHI_Narcissus_tazetta@1
CHI_Asparagus_officinalis@18144
CHI_Phoenix_dactylifera@2205
CHI_Elaeis_guineensis@12604
CHI_Musa_acuminata@12129
CHI_Acomosus_321_v3@7433
CHI_Ananas_comosus@9358
CHI_Lilium_hybrid@1
CHI_Dalata_550_v2_1@9327
CHI_Amborella_trichopoda@10030
0.2

### FLS

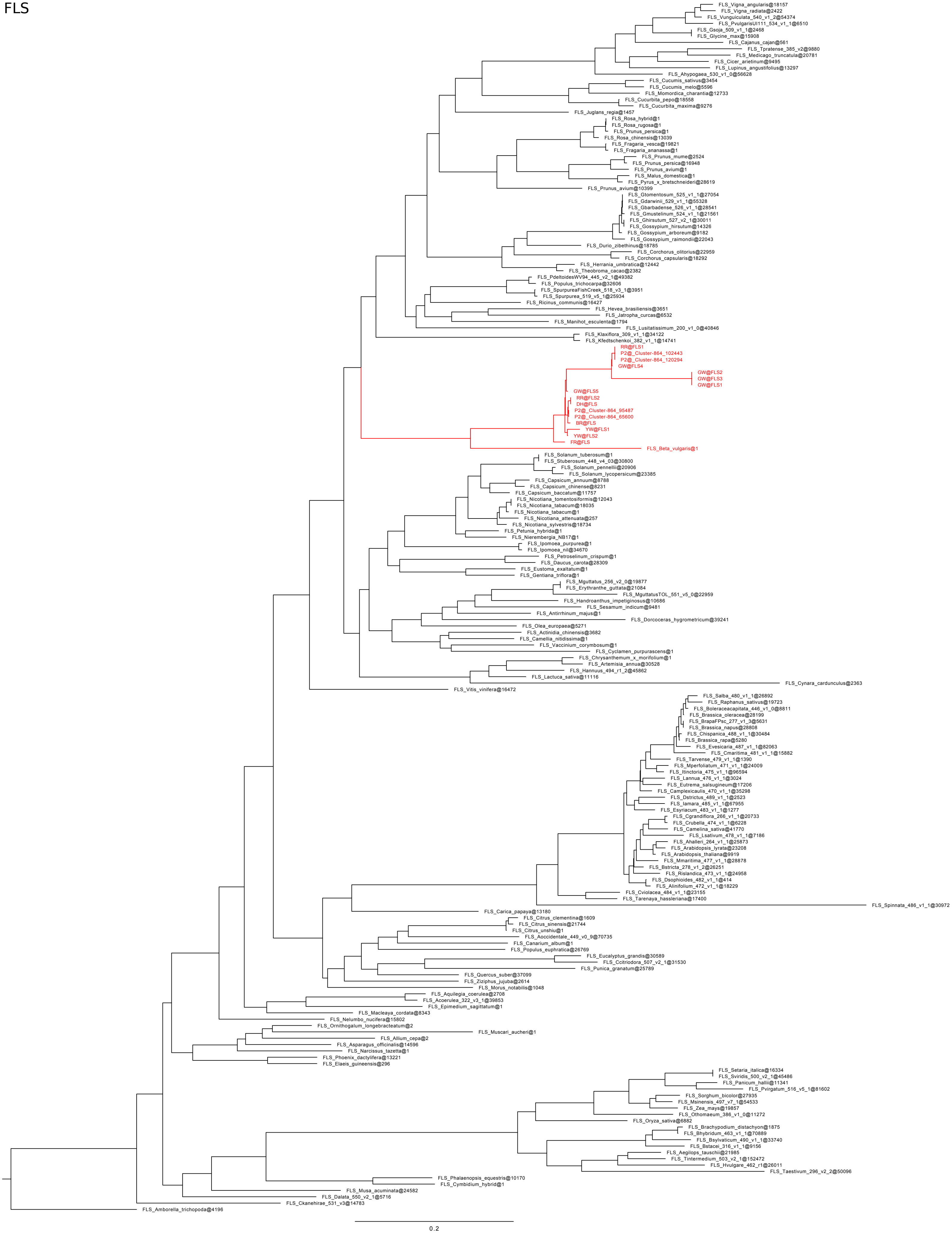
FLS_Vigna_angularis@18157
FLS_Vigna_radiata@2422
FLS_Vunguiculata_540_v1_2@54374
FLS_PvulgarisUI111_534_v1_1@6510
FLS_Gsoja_509_v1_1@2468
FLS_Glycine_max@15908
FLS_Cajanus_cajan@561
FLS_Tpratense_385_v2@9880
FLS_Medicago_truncatula@20781
FLS_Cicer_arietinum@9495
FLS_Lupinus_angustifolius@13297
FLS_Ahypogaea_530_v1_0@56628
FLS_Cucumis_sativus@3454
FLS_Cucumis_melo@5596
FLS_Momordica_charantia@12733
FLS_Cucurbita_pepo@18558
FLS_Cucurbita_maxima@9276
FLS_Juglans_regia@1457
FLS_Rosa_hybrid@1
FLS_Rosa_rugosa@1
FLS_Prunus_persica@1
FLS_Rosa_chinensis@13039
FLS_Fragaria_vesca@19821
FLS_Fragaria_ananassa@1
FLS_Prunus_mume@2524
FLS_Prunus_persica@16948
FLS_Prunus_avium@1
FLS_Malus_domestica@1
FLS_Pyrus_x_bretschneideri@28619
FLS_Prunus_avium@10399
FLS_Gtomentosum_525_v1_1@27054
FLS_Gdarwinii_529_v1_1@55328
FLS_Gbarbadense_526_v1_1@28541
FLS_Gmustelinum_524_v1_1@21561
FLS_Ghirsutum_527_v2_1@30011
FLS_Gossypium_hirsutum@14326
FLS_Gossypium_arboreum@9182
FLS_Gossypium_raimondii@22043
FLS_Durio_zibethinus@18785
FLS_Corchorus_olitorius@22959
FLS_Corchorus_capsularis@18292
FLS_Herrania_umbratica@12442
FLS_Theobroma_cacao@2382
FLS_PdeltoidesWV94_445_v2_1@49382
FLS_Populus_trichocarpa@32606
FLS_SpurpureaFishCreek_518_v3_1@3951
FLS_Spurpurea_519_v5_1@25934
FLS_Ricinus_communis@16427
FLS_Hevea_brasiliensis@3651
FLS_Jatropha_curcas@6532
FLS_Manihot_esculenta@1794
FLS_Lusitatissimum_200_v1_0@40846
FLS_Klaxiflora_309_v1_1@34122
FLS_Kfedtschenkoi_382_v1_1@14741
RR@FLS1
P2@_Cluster-864_102443
P2@_Cluster-864_120294
GW@FLS4
GW@FLS2
GW@FLS3
GW@FLS1
GW@FLS5
RR@FLS2
DH@FLS
P2@_Cluster-864_95487
P2@_Cluster-864_65600
BR@FLS
YW@FLS1
YW@FLS2
FR@FLS
FLS_Beta_vulgaris@1
FLS_Solanum_tuberosum@1
FLS_Stuberosum_448_v4_03@30800
FLS_Solanum_pennellii@20906
FLS_Solanum_lycopersicum@23385
FLS_Capsicum_annuum@8788
FLS_Capsicum_chinense@8231
FLS_Capsicum_baccatum@11757
FLS_Nicotiana_tomentosiformis@12043
FLS_Nicotiana_tabacum@18035
FLS_Nicotiana_tabacum@1
FLS_Nicotiana_attenuata@257
FLS_Nicotiana_sylvestris@18734
FLS_Petunia_hybrida@1
FLS_Nierembergia_NB17@1
FLS_Ipomoea_purpurea@1
FLS_Ipomoea_nil@34670
FLS_Petroselinum_crispum@1
FLS_Daucus_carota@28309
FLS_Eustoma_exaltatum@1
FLS_Gentiana_triflora@1
FLS_Mguttatus_256_v2_0@19877
FLS_Erythranthe_guttata@21084
FLS_MguttatusTOL_551_v5_0@22959
FLS_Handroanthus_impetiginosus@10686
FLS_Sesamum_indicum@9481
FLS_Antirrhinum_majus@1
FLS_Dorcoceras_hygrometricum@39241
FLS_Olea_europaea@5271
FLS_Actinidia_chinensis@3682
FLS_Camellia_nitidissima@1
FLS_Vaccinium_corymbosum@1
FLS_Cyclamen_purpurascens@1
FLS_Chrysanthemum_x_morifolium@1
FLS_Artemisia_annua@30528
FLS_Hannuus_494_r1_2@45862
FLS_Lactuca_sativa@11116
FLS_Cynara_cardunculus@2363
FLS_Vitis_vinifera@16472
FLS_Salba_480_v1_1@26892
FLS_Raphanus_sativus@19723
FLS_Boleraceacapitata_446_v1_0@8811
FLS_Brassica_oleracea@28199
FLS_BrapaFPsc_277_v1_3@5631
FLS_Brassica_napus@28808
FLS_Chispanica_488_v1_1@30484
FLS_Brassica_rapa@5280
FLS_Evesicaria_487_v1_1@82063
FLS_Cmaritima_481_v1_1@15882
FLS_Tarvense_479_v1_1@1390
FLS_Mperfoliatum_471_v1_1@24009
FLS_Itinctoria_475_v1_1@96594
FLS_Lannua_476_v1_1@3024
FLS_Eutrema_salsugineum@17206
FLS_Camplexicaulis_470_v1_1@35298
FLS_Dstrictus_489_v1_1@2523
FLS_Iamara_485_v1_1@67955
FLS_Esyriacum_483_v1_1@1277
FLS_Cgrandiflora_266_v1_1@20733
FLS_Crubella_474_v1_1@6228
FLS_Camelina_sativa@41770
FLS_Lsativum_478_v1_1@7186
FLS_Ahalleri_264_v1_1@25873
FLS_Arabidopsis_lyrata@23208
FLS_Arabidopsis_thaliana@9919
FLS_Mmaritima_477_v1_1@28878
FLS_Bstricta_278_v1_2@26251
FLS_Rislandica_473_v1_1@24958
FLS_Dsophioides_482_v1_1@414
FLS_Alinifolium_472_v1_1@18229
FLS_Cviolacea_484_v1_1@23155
FLS_Tarenaya_hassleriana@17400
FLS_Spinnata_486_v1_1@30972
FLS_Carica_papaya@13180
FLS_Citrus_clementina@1609
FLS_Citrus_sinensis@21744
FLS_Citrus_unshiu@1
FLS_Aoccidentale_449_v0_9@70735
FLS_Canarium_album@1
FLS_Populus_euphratica@26769
FLS_Eucalyptus_grandis@30589
FLS_Ccitriodora_507_v2_1@31530
FLS_Punica_granatum@25789
FLS_Quercus_suber@37099
FLS_Ziziphus_jujuba@2614
FLS_Morus_notabilis@1048
FLS_Aquilegia_coerulea@2708
FLS_Acoerulea_322_v3_1@39853
FLS_Epimedium_sagittatum@1
FLS_Macleaya_cordata@8343
FLS_Nelumbo_nucifera@15802
FLS_Ornithogalum_longebracteatum@2
FLS_Muscari_aucheri@1
FLS_Allium_cepa@2
FLS_Asparagus_officinalis@14596
FLS_Narcissus_tazetta@1
FLS_Phoenix_dactylifera@13221
FLS_Elaeis_guineensis@296
FLS_Setaria_italica@16334
FLS_Sviridis_500_v2_1@45486
FLS_Panicum_hallii@11341
FLS_Pvirgatum_516_v5_1@81602
FLS_Sorghum_bicolor@27935
FLS_Msinensis_497_v7_1@54533
FLS_Zea_mays@19857
FLS_Othomaeum_386_v1_0@11272
FLS_Oryza_sativa@6882
FLS_Brachypodium_distachyon@1875
FLS_Bhybridum_463_v1_1@70889
FLS_Bsylvaticum_490_v1_1@33740
FLS_Bstacei_316_v1_1@9156
FLS_Aegilops_tauschii@21985
FLS_Tintermedium_503_v2_1@152472
FLS_Hvulgare_462_r1@26011
FLS_Taestivum_296_v2_2@50096
FLS_Phalaenopsis_equestris@10170
FLS_Cymbidium_hybrid@1
FLS_Musa_acuminata@24582
FLS_Dalata_550_v2_1@5716
FLS_Ckanehirae_531_v3@14783
FLS_Amborella_trichopoda@4196
0.2

F3H

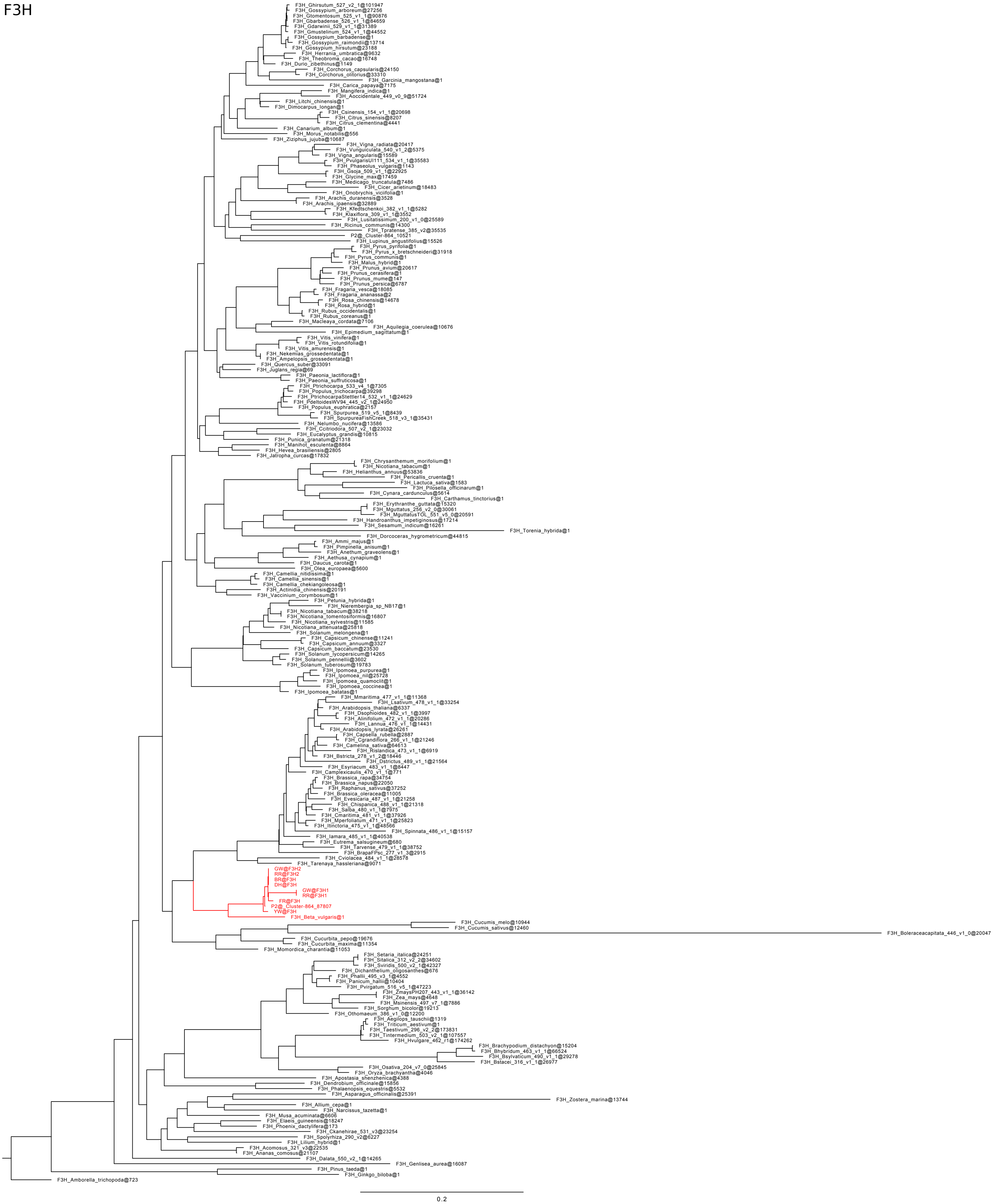

F3H_Ghirsutum_527_v2_1@101947
F3H_Gossypium_arboreum@27256
F3H_Gtomentosum_525_v1_1@90876
F3H_Gbarbadense_526_v1_1@84659
F3H_Gdarwinii_529_v1_1@31389
F3H_Gmustelinum_524_v1_1@44552
F3H_Gossypium_barbadense@1
F3H_Gossypium_raimondii@13714
F3H_Gossypium_hirsutum@23188
F3H_Herrania_umbratica@9632
F3H_Theobroma_cacao@16748
F3H_Durio_zibethinus@1149
F3H_Corchorus_capsularis@24150
F3H_Corchorus_olitorius@33310
F3H_Garcinia_mangostana@1
F3H_Carica_papaya@7175
F3H_Mangifera_indica@1
F3H_Aoccidentale_449_v0_9@51724
F3H_Litchi_chinensis@1
F3H_Dimocarpus_longan@1
F3H_Csinensis_154_v1_1@20698
F3H_Citrus_sinensis@8207
F3H_Citrus_clementina@4441
F3H_Canarium_album@1
F3H_Morus_notabilis@556
F3H_Ziziphus_jujuba@10687
F3H_Vigna_radiata@20417
F3H_Vunguiculata_540_v1_2@5375
F3H_Vigna_angularis@15589
F3H_PvulgarisUI111_534_v1_1@35583
F3H_Phaseolus_vulgaris@1143
F3H_Gsoja_509_v1_1@22925
F3H_Glycine_max@17459
F3H_Medicago_truncatula@7486
F3H_Cicer_arietinum@18483
F3H_Onobrychis_viciifolia@1
F3H_Arachis_duranensis@3528
F3H_Arachis_ipaensis@32889
F3H_Kfedtschenkoi_382_v1_1@5282
F3H_Klaxiflora_309_v1_1@3552
F3H_Lusitatissimum_200_v1_0@25589
F3H_Ricinus_communis@14300
F3H_Tpratense_385_v2@35535
P2@_Cluster-864_10521
F3H_Lupinus_angustifolius@15526
F3H_Pyrus_pyrifolia@1
F3H_Pyrus_x_bretschneideri@31918
F3H_Pyrus_communis@1
F3H_Malus_hybrid@1
F3H_Prunus_avium@20617
F3H_Prunus_cerasifera@1
F3H_Prunus_mume@147
F3H_Prunus_persica@6787
F3H_Fragaria_vesca@18085
F3H_Fragaria_ananassa@2
F3H_Rosa_chinensis@14678
F3H_Rosa_hybrid@1
F3H_Rubus_occidentalis@1
F3H_Rubus_coreanus@1
F3H_Macleaya_cordata@7106
F3H_Aquilegia_coerulea@10676
F3H_Epimedium_sagittatum@1
F3H_Vitis_vinifera@1
F3H_Vitis_rotundifolia@1
F3H_Vitis_amurensis@1
F3H_Nekemias_grossedentata@1
F3H_Ampelopsis_grossedentata@1
F3H_Quercus_suber@33091
F3H_Juglans_regia@69
F3H_Paeonia_lactiflora@1
F3H_Paeonia_suffruticosa@1
F3H_Ptrichocarpa_533_v4_1@7305
F3H_Populus_trichocarpa@39298
F3H_PtrichocarpaStettler14_532_v1_1@24629
F3H_PdeltoidesWV94_445_v2_1@24950
F3H_Populus_euphratica@2157
F3H_Spurpurea_519_v5_1@8439
F3H_SpurpureaFishCreek_518_v3_1@35431
F3H_Nelumbo_nucifera@13586
F3H_Ccitriodora_507_v2_1@23032
F3H_Eucalyptus_grandis@10815
F3H_Punica_granatum@21318
F3H_Manihot_esculenta@8864
F3H_Hevea_brasiliensis@2805
F3H_Jatropha_curcas@17832
F3H_Chrysanthemum_morifolium@1
F3H_Nicotiana_tabacum@1
F3H_Helianthus_annuus@53836
F3H_Pericallis_cruenta@1
F3H_Lactuca_sativa@1583
F3H_Pilosella_officinarum@1
F3H_Cynara_cardunculus@5614
F3H_Carthamus_tinctorius@1
F3H_Erythranthe_guttata@15320
F3H_Mguttatus_256_v2_0@30061
F3H_MguttatusTOL_551_v5_0@20591
F3H_Handroanthus_impetiginosus@17214
F3H_Sesamum_indicum@16261
F3H_Torenia_hybrida@1
F3H_Dorcoceras_hygrometricum@44815
F3H_Ammi_majus@1
F3H_Pimpinella_anisum@1
F3H_Anethum_graveolens@1
F3H_Aethusa_cynapium@1
F3H_Daucus_carota@1
F3H_Olea_europaea@5600
F3H_Camellia_nitidissima@1
F3H_Camellia_sinensis@1
F3H_Camellia_chekiangoleosa@1
F3H_Actinidia_chinensis@20191
F3H_Vaccinium_corymbosum@1
F3H_Petunia_hybrida@1
F3H_Nierembergia_sp_NB17@1
F3H_Nicotiana_tabacum@38218
F3H_Nicotiana_tomentosiformis@16807
F3H_Nicotiana_sylvestris@11585
F3H_Nicotiana_attenuata@25818
F3H_Solanum_melongena@1
F3H_Capsicum_chinense@11241
F3H_Capsicum_annuum@3327
F3H_Capsicum_baccatum@23530
F3H_Solanum_lycopersicum@14265
F3H_Solanum_pennellii@3602
F3H_Solanum_tuberosum@19783
F3H_Ipomoea_purpurea@1
F3H_Ipomoea_nil@25728
F3H_Ipomoea_quamoclit@1
F3H_Ipomoea_coccinea@1
F3H_Ipomoea_batatas@1
F3H_Mmaritima_477_v1_1@11368
F3H_Lsativum_478_v1_1@33254
F3H_Arabidopsis_thaliana@6337
F3H_Dsophioides_482_v1_1@3997
F3H_Alinifolium_472_v1_1@20286
F3H_Lannua_476_v1_1@14431
F3H_Arabidopsis_lyrata@26261
F3H_Capsella_rubella@2887
F3H_Cgrandiflora_266_v1_1@21246
F3H_Camelina_sativa@64613
F3H_Rislandica_473_v1_1@6919
F3H_Bstricta_278_v1_2@18446
F3H_Dstrictus_489_v1_1@21564
F3H_Esyriacum_483_v1_1@8447
F3H_Camplexicaulis_470_v1_1@771
F3H_Brassica_rapa@34754
F3H_Brassica_napus@22050
F3H_Raphanus_sativus@37252
F3H_Brassica_oleracea@11005
F3H_Evesicaria_487_v1_1@21258
F3H_Chispanica_488_v1_1@21318
F3H_Salba_480_v1_1@7975
F3H_Cmaritima_481_v1_1@37926
F3H_Mperfoliatum_471_v1_1@25823
F3H_Itinctoria_475_v1_1@48566
F3H_Spinnata_486_v1_1@15157
F3H_Iamara_485_v1_1@40538
F3H_Eutrema_salsugineum@680
F3H_Tarvense_479_v1_1@38752
F3H_BrapaFPsc_277_v1_3@2915
F3H_Cviolacea_484_v1_1@28578
F3H_Tarenaya_hassleriana@9071
GW@F3H2
RR@F3H2
BR@F3H
DH@F3H
GW@F3H1
RR@F3H1
FR@F3H
P2@_Cluster-864_87807
YW@F3H
F3H_Beta_vulgaris@1
F3H_Cucumis_melo@10944
F3H_Cucumis_sativus@12460
F3H_Boleraceacapitata_446_v1_0@20047
F3H_Cucurbita_pepo@19676
F3H_Cucurbita_maxima@11354
F3H_Momordica_charantia@11053
F3H_Setaria_italica@24251
F3H_Sitalica_312_v2_2@34602
F3H_Sviridis_500_v2_1@42327
F3H_Dichanthelium_oligosanthes@676
F3H_Phallii_495_v3_1@4552
F3H_Panicum_hallii@10404
F3H_Pvirgatum_516_v5_1@47223
F3H_ZmaysPH207_443_v1_1@36142
F3H_Zea_mays@4648
F3H_Msinensis_497_v7_1@7886
F3H_Sorghum_bicolor@19213
F3H_Othomaeum_386_v1_0@12200
F3H_Aegilops_tauschii@1319
F3H_Triticum_aestivum@1
F3H_Taestivum_296_v2_2@173831
F3H_Tintermedium_503_v2_1@107557
F3H_Hvulgare_462_r1@174262
F3H_Brachypodium_distachyon@15204
F3H_Bhybridum_463_v1_1@66524
F3H_Bsylvaticum_490_v1_1@29278
F3H_Bstacei_316_v1_1@26977
F3H_Osativa_204_v7_0@25845
F3H_Oryza_brachyantha@4046
F3H_Apostasia_shenzhenica@4388
F3H_Dendrobium_officinale@15856
F3H_Phalaenopsis_equestris@5532
F3H_Asparagus_officinalis@25391
F3H_Zostera_marina@13744
F3H_Allium_cepa@1
F3H_Narcissus_tazetta@1
F3H_Musa_acuminata@6606
F3H_Elaeis_guineensis@18247
F3H_Phoenix_dactylifera@173
F3H_Ckanehirae_531_v3@23254
F3H_Spolyrhiza_290_v2@6227
F3H_Lilium_hybrid@1
F3H_Acomosus_321_v3@22535
F3H_Ananas_comosus@21107
F3H_Dalata_550_v2_1@14265
F3H_Genlisea_aurea@16087
F3H_Pinus_taeda@1
F3H_Ginkgo_biloba@1
F3H_Amborella_trichopoda@723
0.2

F3'H

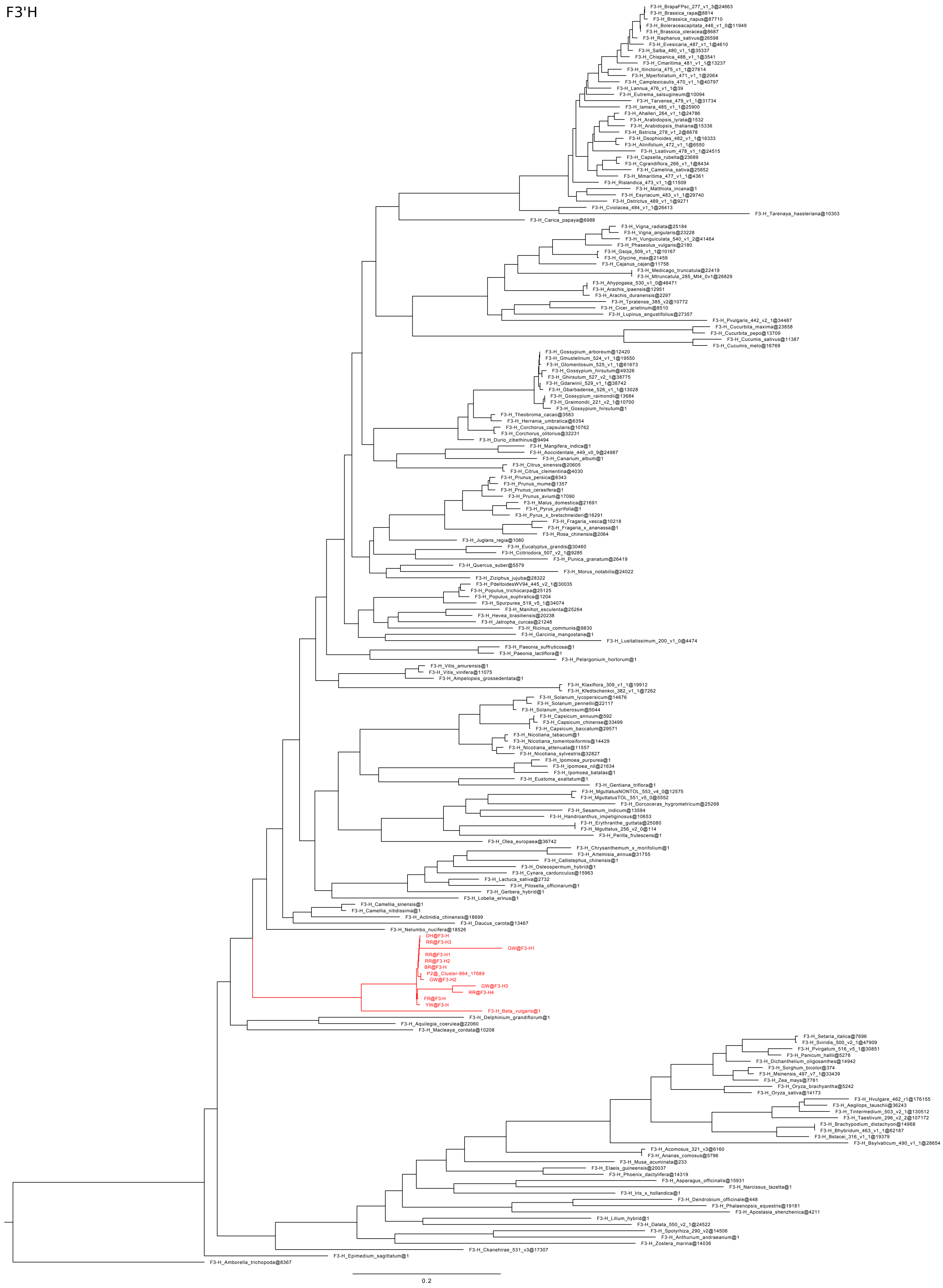
F3-H_BrapaFPsc_277_v1_3@24663
F3-H_Brassica_rapa@8814
F3-H_Brassica_napus@87710
F3-H_Boleraceacapitata_446_v1_0@11949
F3-H_Brassica_oleracea@8687
F3-H_Raphanus_sativus@26598
F3-H_Evesicaria_487_v1_1@4610
F3-H_Salba_480_v1_1@35337
F3-H_Chispanica_488_v1_1@3541
F3-H_Cmaritima_481_v1_1@13237
F3-H_Itinctoria_475_v1_1@27614
F3-H_Mperfoliatum_471_v1_1@2064
F3-H_Camplexicaulis_470_v1_1@40797
F3-H_Lannua_476_v1_1@39
F3-H_Eutrema_salsugineum@10094
F3-H_Tarvense_479_v1_1@31734
F3-H_Iamara_485_v1_1@25900
F3-H_Ahalleri_264_v1_1@24786
F3-H_Arabidopsis_lyrata@1532
F3-H_Arabidopsis_thaliana@15336
F3-H_Bstricta_278_v1_2@8678
F3-H_Dsophioides_482_v1_1@16333
F3-H_Alinifolium_472_v1_1@6550
F3-H_Lsativum_478_v1_1@24515
F3-H_Capsella_rubella@23689
F3-H_Cgrandiflora_266_v1_1@8434
F3-H_Camelina_sativa@25852
F3-H_Mmaritima_477_v1_1@4361
F3-H_Rislandica_473_v1_1@11509
F3-H_Matthiola_incana@1
F3-H_Esyriacum_483_v1_1@29740
F3-H_Dstrictus_489_v1_1@9271
F3-H_Cviolacea_484_v1_1@26413
F3-H_Tarenaya_hassleriana@10303
F3-H_Carica_papaya@6988
F3-H_Vigna_radiata@25184
F3-H_Vigna_angularis@23228
F3-H_Vunguiculata_540_v1_2@41464
F3-H_Phaseolus_vulgaris@2180
F3-H_Gsoja_509_v1_1@10167
F3-H_Glycine_max@21459
F3-H_Cajanus_cajan@11758
F3-H_Medicago_truncatula@22419
F3-H_Mtruncatula_285_Mt4_0v1@26829
F3-H_Ahypogaea_530_v1_0@46471
F3-H_Arachis_ipaensis@12951
F3-H_Arachis_duranensis@2297
F3-H_Tpratense_385_v2@10772
F3-H_Cicer_arietinum@8510
F3-H_Lupinus_angustifolius@27357
F3-H_Pvulgaris_442_v2_1@34487
F3-H_Cucurbita_maxima@23858
F3-H_Cucurbita_pepo@13709
F3-H_Cucumis_sativus@11387
F3-H_Cucumis_melo@16769
F3-H_Gossypium_arboreum@12420
F3-H_Gmustelinum_524_v1_1@19550
F3-H_Gtomentosum_525_v1_1@61673
F3-H_Gossypium_hirsutum@49326
F3-H_Ghirsutum_527_v2_1@38775
F3-H_Gdarwinii_529_v1_1@38742
F3-H_Gbarbadense_526_v1_1@13028
F3-H_Gossypium_raimondii@13684
F3-H_Graimondii_221_v2_1@10700
F3-H_Gossypium_hirsutum@1
F3-H_Theobroma_cacao@3583
F3-H_Herrania_umbratica@6354
F3-H_Corchorus_capsularis@10762
F3-H_Corchorus_olitorius@32231
F3-H_Durio_zibethinus@9494
F3-H_Mangifera_indica@1
F3-H_Aoccidentale_449_v0_9@24987
F3-H_Canarium_album@1
F3-H_Citrus_sinensis@20605
F3-H_Citrus_clementina@4030
F3-H_Prunus_persica@8343
F3-H_Prunus_mume@1357
F3-H_Prunus_cerasifera@1
F3-H_Prunus_avium@17090
F3-H_Malus_domestica@21691
F3-H_Pyrus_pyrifolia@1
F3-H_Pyrus_x_bretschneideri@16291
F3-H_Fragaria_vesca@10218
F3-H_Fragaria_x_ananassa@1
F3-H_Rosa_chinensis@2064
F3-H_Juglans_regia@1080
F3-H_Eucalyptus_grandis@30460
F3-H_Ccitriodora_507_v2_1@9285
F3-H_Punica_granatum@26419
F3-H_Quercus_suber@5579
F3-H_Morus_notabilis@24022
F3-H_Ziziphus_jujuba@28322
F3-H_PdeltoidesWV94_445_v2_1@30035
F3-H_Populus_trichocarpa@25125
F3-H_Populus_euphratica@1204
F3-H_Spurpurea_519_v5_1@34074
F3-H_Manihot_esculenta@25264
F3-H_Hevea_brasiliensis@20238
F3-H_Jatropha_curcas@21248
F3-H_Ricinus_communis@9830
F3-H_Garcinia_mangostana@1
F3-H_Lusitatissimum_200_v1_0@4474
F3-H_Paeonia_suffruticosa@1
F3-H_Paeonia_lactiflora@1
F3-H_Pelargonium_hortorum@1
F3-H_Vitis_amurensis@1
F3-H_Vitis_vinifera@11075
F3-H_Ampelopsis_grossedentata@1
F3-H_Klaxiflora_309_v1_1@19912
F3-H_Kfedtschenkoi_382_v1_1@7262
F3-H_Solanum_lycopersicum@14676
F3-H_Solanum_pennellii@22117
F3-H_Solanum_tuberosum@5044
F3-H_Capsicum_annuum@592
F3-H_Capsicum_chinense@33499
F3-H_Capsicum_baccatum@29571
F3-H_Nicotiana_tabacum@1
F3-H_Nicotiana_tomentosiformis@14429
F3-H_Nicotiana_attenuata@11557
F3-H_Nicotiana_sylvestris@32827
F3-H_Ipomoea_purpurea@1
F3-H_Ipomoea_nil@21634
F3-H_Ipomoea_batatas@1
F3-H_Eustoma_exaltatum@1
F3-H_Gentiana_triflora@1
F3-H_MguttatusNONTOL_553_v4_0@12575
F3-H_MguttatusTOL_551_v5_0@5552
F3-H_Dorcoceras_hygrometricum@25268
F3-H_Sesamum_indicum@13594
F3-H_Handroanthus_impetiginosus@10653
F3-H_Erythranthe_guttata@25080
F3-H_Mguttatus_256_v2_0@114
F3-H_Perilla_frutescens@1
F3-H_Olea_europaea@36742
F3-H_Chrysanthemum_x_morifolium@1
F3-H_Artemisia_annua@31755
F3-H_Callistephus_chinensis@1
F3-H_Osteospermum_hybrid@1
F3-H_Cynara_cardunculus@15963
F3-H_Lactuca_sativa@2732
F3-H_Pilosella_officinarum@1
F3-H_Gerbera_hybrid@1
F3-H_Lobelia_erinus@1
F3-H_Camellia_sinensis@1
F3-H_Camellia_nitidissima@1
F3-H_Actinidia_chinensis@18699
F3-H_Daucus_carota@13467
F3-H_Nelumbo_nucifera@18526
DH@F3-H
RR@F3-H3
GW@F3-H1
RR@F3-H1
RR@F3-H2
BR@F3-H
P2@_Cluster-864_17689
GW@F3-H2
GW@F3-H3
RR@F3-H4
FR@F3-H
YW@F3-H
F3-H_Beta_vulgaris@1
F3-H_Delphinium_grandiflorum@1
F3-H_Aquilegia_coerulea@22060
F3-H_Macleaya_cordata@10208
F3-H_Setaria_italica@7696
F3-H_Sviridis_500_v2_1@47909
F3-H_Pvirgatum_516_v5_1@30851
F3-H_Panicum_hallii@5278
F3-H_Dichanthelium_oligosanthes@14942
F3-H_Sorghum_bicolor@374
F3-H_Msinensis_497_v7_1@33439
F3-H_Zea_mays@7781
F3-H_Oryza_brachyantha@5242
F3-H_Oryza_sativa@14173
F3-H_Hvulgare_462_r1@176155
F3-H_Aegilops_tauschii@36243
F3-H_Tintermedium_503_v2_1@130512
F3-H_Taestivum_296_v2_2@107172
F3-H_Brachypodium_distachyon@14968
F3-H_Bhybridum_463_v1_1@62187
F3-H_Bstacei_316_v1_1@19379
F3-H_Bsylvaticum_490_v1_1@28654
F3-H_Acomosus_321_v3@6160
F3-H_Ananas_comosus@5796
F3-H_Musa_acuminata@233
F3-H_Elaeis_guineensis@20037
F3-H_Phoenix_dactylifera@14319
F3-H_Asparagus_officinalis@15931
F3-H_Narcissus_tazetta@1
F3-H_Iris_x_hollandica@1
F3-H_Dendrobium_officinale@448
F3-H_Phalaenopsis_equestris@19181
F3-H_Apostasia_shenzhenica@4211
F3-H_Lilium_hybrid@1
F3-H_Dalata_550_v2_1@24522
F3-H_Spolyrhiza_290_v2@14506
F3-H_Anthurium_andraeanum@1
F3-H_Zostera_marina@14036
F3-H_Ckanehirae_531_v3@17307
F3-H_Epimedium_sagittatum@1
F3-H_Amborella_trichopoda@8367
0.2

### DFR

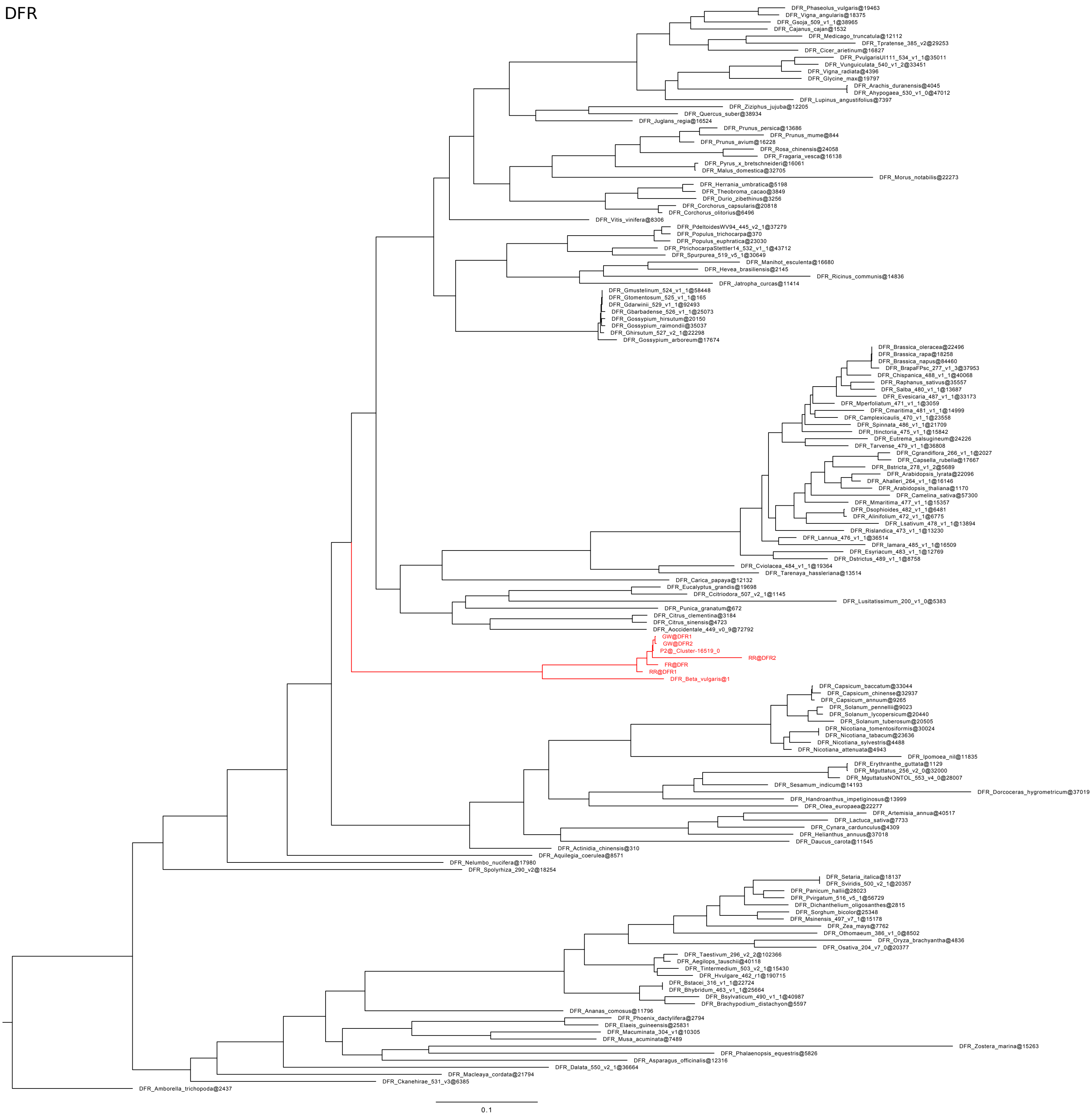
DFR_Phaseolus_vulgaris@19463
DFR_Vigna_angularis@18375
DFR_Gsoja_509_v1_1@38965
DFR_Cajanus_cajan@1532
DFR_Medicago_truncatula@12112
DFR_Tpratense_385_v2@29253
DFR_Cicer_arietinum@16827
DFR_PvulgarisUI111_534_v1_1@35011
DFR_Vunguiculata_540_v1_2@33451
DFR_Vigna_radiata@4396
DFR_Glycine_max@19797
DFR_Arachis_duranensis@4045
DFR_Ahypogaea_530_v1_0@47012
DFR_Lupinus_angustifolius@7397
DFR_Ziziphus_jujuba@12205
DFR_Quercus_suber@38934
DFR_Juglans_regia@16524
DFR_Prunus_persica@13686
DFR_Prunus_mume@844
DFR_Prunus_avium@16228
DFR_Rosa_chinensis@24058
DFR_Fragaria_vesca@16138
DFR_Pyrus_x_bretschneideri@16061
DFR_Malus_domestica@32705
DFR_Morus_notabilis@22273
DFR_Herrania_umbratica@5198
DFR_Theobroma_cacao@3849
DFR_Durio_zibethinus@3256
DFR_Corchorus_capsularis@20818
DFR_Corchorus_olitorius@6496
DFR_Vitis_vinifera@8306
DFR_PdeltoidesWV94_445_v2_1@37279
DFR_Populus_trichocarpa@370
DFR_Populus_euphratica@23030
DFR_PtrichocarpaStettler14_532_v1_1@43712
DFR_Spurpurea_519_v5_1@30649
DFR_Manihot_esculenta@16680
DFR_Hevea_brasiliensis@2145
DFR_Ricinus_communis@14836
DFR_Jatropha_curcas@11414
DFR_Gmustelinum_524_v1_1@58448
DFR_Gtomentosum_525_v1_1@165
DFR_Gdarwinii_529_v1_1@92493
DFR_Gbarbadense_526_v1_1@25073
DFR_Gossypium_hirsutum@20150
DFR_Gossypium_raimondii@35037
DFR_Ghirsutum_527_v2_1@22298
DFR_Gossypium_arboreum@17674
DFR_Brassica_oleracea@22496
DFR_Brassica_rapa@18258
DFR_Brassica_napus@84460
DFR_BrapaFPsc_277_v1_3@37953
DFR_Chispanica_488_v1_1@40068
DFR_Raphanus_sativus@35557
DFR_Salba_480_v1_1@13687
DFR_Evesicaria_487_v1_1@33173
DFR_Mperfoliatum_471_v1_1@3059
DFR_Cmaritima_481_v1_1@14999
DFR_Camplexicaulis_470_v1_1@23558
DFR_Spinnata_486_v1_1@21709
DFR_Itinctoria_475_v1_1@15842
DFR_Eutrema_salsugineum@24226
DFR_Tarvense_479_v1_1@36808
DFR_Cgrandiflora_266_v1_1@2027
DFR_Capsella_rubella@17667
DFR_Bstricta_278_v1_2@5689
DFR_Arabidopsis_lyrata@22096
DFR_Ahalleri_264_v1_1@16146
DFR_Arabidopsis_thaliana@1170
DFR_Camelina_sativa@57300
DFR_Mmaritima_477_v1_1@15357
DFR_Dsophioides_482_v1_1@6481
DFR_Alinifolium_472_v1_1@6775
DFR_Lsativum_478_v1_1@13894
DFR_Rislandica_473_v1_1@13230
DFR_Lannua_476_v1_1@36514
DFR_Iamara_485_v1_1@16509
DFR_Esyriacum_483_v1_1@12769
DFR_Dstrictus_489_v1_1@8758
DFR_Cviolacea_484_v1_1@19364
DFR_Tarenaya_hassleriana@13514
DFR_Carica_papaya@12132
DFR_Eucalyptus_grandis@19698
DFR_Ccitriodora_507_v2_1@1145
DFR_Lusitatissimum_200_v1_0@5383
DFR_Punica_granatum@672
DFR_Citrus_clementina@3184
DFR_Citrus_sinensis@4723
DFR_Aoccidentale_449_v0_9@72792
GW@DFR1
GW@DFR2
P2@_Cluster-16519_0
RR@DFR2
FR@DFR
RR@DFR1
DFR_Beta_vulgaris@1
DFR_Capsicum_baccatum@33044
DFR_Capsicum_chinense@32937
DFR_Capsicum_annuum@9265
DFR_Solanum_pennellii@9023
DFR_Solanum_lycopersicum@20440
DFR_Solanum_tuberosum@20505
DFR_Nicotiana_tomentosiformis@30024
DFR_Nicotiana_tabacum@23636
DFR_Nicotiana_sylvestris@4488
DFR_Nicotiana_attenuata@4943
DFR_Ipomoea_nil@11835
DFR_Erythranthe_guttata@1129
DFR_Mguttatus_256_v2_0@32000
DFR_MguttatusNONTOL_553_v4_0@28007
DFR_Sesamum_indicum@14193
DFR_Dorcoceras_hygrometricum@37019
DFR_Handroanthus_impetiginosus@13999
DFR_Olea_europaea@22277
DFR_Artemisia_annua@40517
DFR_Lactuca_sativa@7733
DFR_Cynara_cardunculus@4309
DFR_Helianthus_annuus@37018
DFR_Daucus_carota@11545
DFR_Actinidia_chinensis@310
DFR_Aquilegia_coerulea@8571
DFR_Nelumbo_nucifera@17980
DFR_Spolyrhiza_290_v2@18254
DFR_Setaria_italica@18137
DFR_Sviridis_500_v2_1@20357
DFR_Panicum_hallii@28023
DFR_Pvirgatum_516_v5_1@56729
DFR_Dichanthelium_oligosanthes@2815
DFR_Sorghum_bicolor@25348
DFR_Msinensis_497_v7_1@15178
DFR_Zea_mays@7762
DFR_Othomaeum_386_v1_0@8502
DFR_Oryza_brachyantha@4836
DFR_Osativa_204_v7_0@20377
DFR_Taestivum_296_v2_2@102366
DFR_Aegilops_tauschii@40118
DFR_Tintermedium_503_v2_1@15430
DFR_Hvulgare_462_r1@190715
DFR_Bstacei_316_v1_1@22724
DFR_Bhybridum_463_v1_1@25664
DFR_Bsylvaticum_490_v1_1@40987
DFR_Brachypodium_distachyon@5597
DFR_Ananas_comosus@11796
DFR_Phoenix_dactylifera@2794
DFR_Elaeis_guineensis@25831
DFR_Macuminata_304_v1@10305
DFR_Musa_acuminata@7489
DFR_Zostera_marina@15263
DFR_Phalaenopsis_equestris@5826
DFR_Asparagus_officinalis@12316
DFR_Dalata_550_v2_1@36664
DFR_Macleaya_cordata@21794
DFR_Ckanehirae_531_v3@6385
DFR_Amborella_trichopoda@2437
0.1

ANS

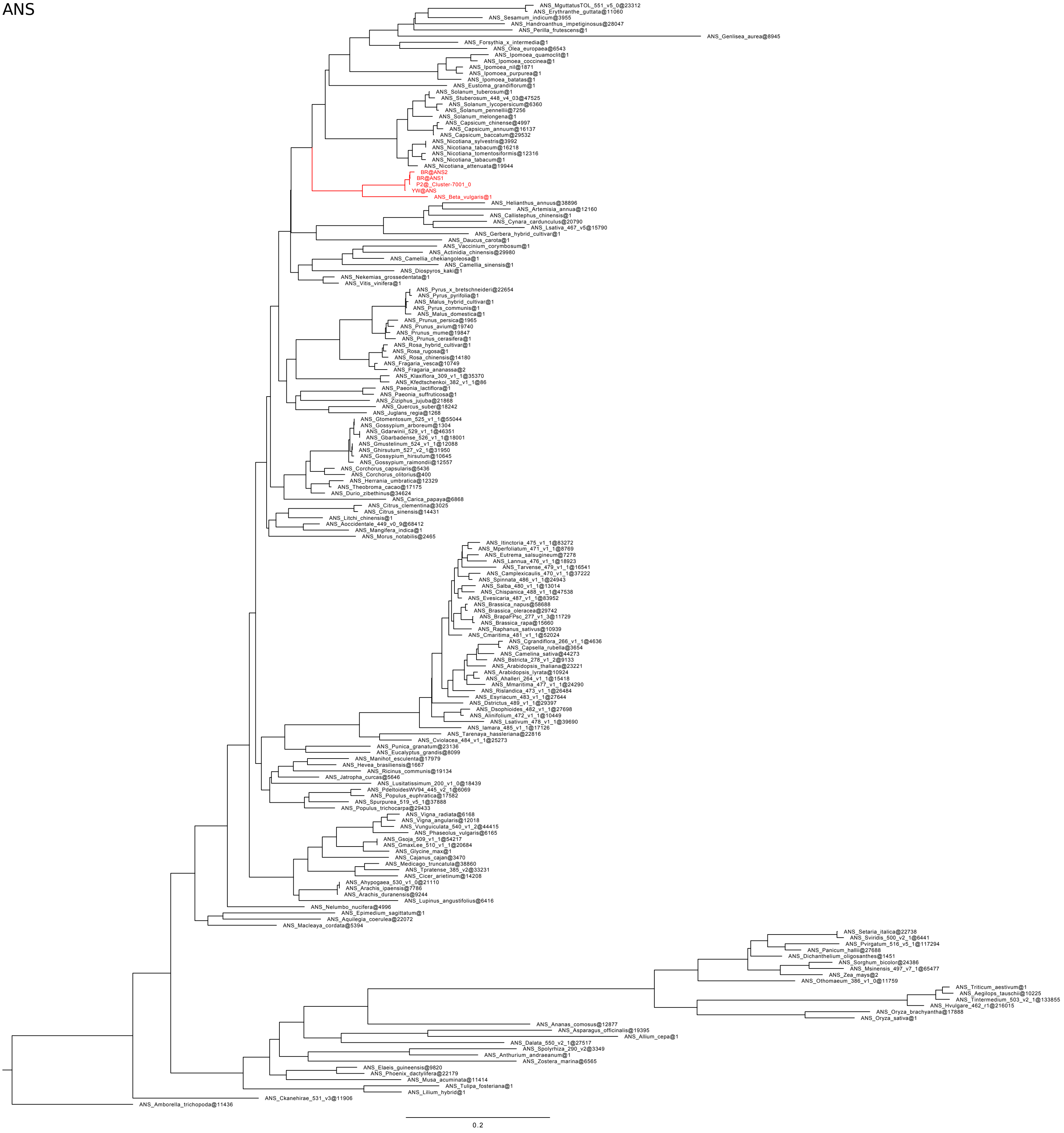
ANS_MguttatusTOL_551_v5_0@23312
ANS_Erythranthe_guttata@11060
ANS_Sesamum_indicum@3955
ANS_Handroanthus_impetiginosus@28047
ANS_Perilla_frutescens@1
ANS_Genlisea_aurea@8945
ANS_Forsythia_x_intermedia@1
ANS_Olea_europaea@6543
ANS_Ipomoea_quamoclit@1
ANS_Ipomoea_coccinea@1
ANS_Ipomoea_nil@1871
ANS_Ipomoea_purpurea@1
ANS_Ipomoea_batatas@1
ANS_Eustoma_grandiflorum@1
ANS_Solanum_tuberosum@1
ANS_Stuberosum_448_v4_03@47525
ANS_Solanum_lycopersicum@6360
ANS_Solanum_pennellii@7256
ANS_Solanum_melongena@1
ANS_Capsicum_chinense@4997
ANS_Capsicum_annuum@16137
ANS_Capsicum_baccatum@29532
ANS_Nicotiana_sylvestris@3992
ANS_Nicotiana_tabacum@16218
ANS_Nicotiana_tomentosiformis@12316
ANS_Nicotiana_tabacum@1
ANS_Nicotiana_attenuata@19944
BR@ANS2
BR@ANS1
P2@_Cluster-7001_0
YW@ANS
ANS_Beta_vulgaris@1
ANS_Helianthus_annuus@38896
ANS_Artemisia_annua@12160
ANS_Callistephus_chinensis@1
ANS_Cynara_cardunculus@20790
ANS_Lsativa_467_v5@15790
ANS_Gerbera_hybrid_cultivar@1
ANS_Daucus_carota@1
ANS_Vaccinium_corymbosum@1
ANS_Actinidia_chinensis@29980
ANS_Camellia_chekiangoleosa@1
ANS_Camellia_sinensis@1
ANS_Diospyros_kaki@1
ANS_Nekemias_grossedentata@1
ANS_Vitis_vinifera@1
ANS_Pyrus_x_bretschneideri@22654
ANS_Pyrus_pyrifolia@1
ANS_Malus_hybrid_cultivar@1
ANS_Pyrus_communis@1
ANS_Malus_domestica@1
ANS_Prunus_persica@1965
ANS_Prunus_avium@19740
ANS_Prunus_mume@19847
ANS_Prunus_cerasifera@1
ANS_Rosa_hybrid_cultivar@1
ANS_Rosa_rugosa@1
ANS_Rosa_chinensis@14180
ANS_Fragaria_vesca@10749
ANS_Fragaria_ananassa@2
ANS_Klaxiflora_309_v1_1@35370
ANS_Kfedtschenkoi_382_v1_1@86
ANS_Paeonia_lactiflora@1
ANS_Paeonia_suffruticosa@1
ANS_Ziziphus_jujuba@21868
ANS_Quercus_suber@18242
ANS_Juglans_regia@1268
ANS_Gtomentosum_525_v1_1@55044
ANS_Gossypium_arboreum@1304
ANS_Gdarwinii_529_v1_1@46351
ANS_Gbarbadense_526_v1_1@18001
ANS_Gmustelinum_524_v1_1@12088
ANS_Ghirsutum_527_v2_1@31950
ANS_Gossypium_hirsutum@10645
ANS_Gossypium_raimondii@12557
ANS_Corchorus_capsularis@5436
ANS_Corchorus_olitorius@400
ANS_Herrania_umbratica@12329
ANS_Theobroma_cacao@17175
ANS_Durio_zibethinus@34624
ANS_Carica_papaya@6868
ANS_Citrus_clementina@3025
ANS_Citrus_sinensis@14431
ANS_Litchi_chinensis@1
ANS_Aoccidentale_449_v0_9@68412
ANS_Mangifera_indica@1
ANS_Morus_notabilis@2465
ANS_Itinctoria_475_v1_1@83272
ANS_Mperfoliatum_471_v1_1@8769
ANS_Eutrema_salsugineum@7278
ANS_Lannua_476_v1_1@18923
ANS_Tarvense_479_v1_1@16541
ANS_Camplexicaulis_470_v1_1@37222
ANS_Spinnata_486_v1_1@24943
ANS_Salba_480_v1_1@13014
ANS_Chispanica_488_v1_1@47538
ANS_Evesicaria_487_v1_1@83952
ANS_Brassica_napus@58688
ANS_Brassica_oleracea@29742
ANS_BrapaFPsc_277_v1_3@11729
ANS_Brassica_rapa@15660
ANS_Raphanus_sativus@10939
ANS_Cmaritima_481_v1_1@52024
ANS_Cgrandiflora_266_v1_1@4636
ANS_Capsella_rubella@3654
ANS_Camelina_sativa@44273
ANS_Bstricta_278_v1_2@9133
ANS_Arabidopsis_thaliana@23221
ANS_Arabidopsis_lyrata@10924
ANS_Ahalleri_264_v1_1@15418
ANS_Mmaritima_477_v1_1@24290
ANS_Rislandica_473_v1_1@26484
ANS_Esyriacum_483_v1_1@27644
ANS_Dstrictus_489_v1_1@29397
ANS_Dsophioides_482_v1_1@27698
ANS_Alinifolium_472_v1_1@10449
ANS_Lsativum_478_v1_1@39690
ANS_Iamara_485_v1_1@17126
ANS_Tarenaya_hassleriana@22816
ANS_Cviolacea_484_v1_1@25273
ANS_Punica_granatum@23136
ANS_Eucalyptus_grandis@8099
ANS_Manihot_esculenta@17979
ANS_Hevea_brasiliensis@1667
ANS_Ricinus_communis@19134
ANS_Jatropha_curcas@5646
ANS_Lusitatissimum_200_v1_0@18439
ANS_PdeltoidesWV94_445_v2_1@6069
ANS_Populus_euphratica@17582
ANS_Spurpurea_519_v5_1@37888
ANS_Populus_trichocarpa@29433
ANS_Vigna_radiata@6168
ANS_Vigna_angularis@12018
ANS_Vunguiculata_540_v1_2@44415
ANS_Phaseolus_vulgaris@6165
ANS_Gsoja_509_v1_1@54217
ANS_GmaxLee_510_v1_1@20684
ANS_Glycine_max@1
ANS_Cajanus_cajan@3470
ANS_Medicago_truncatula@38860
ANS_Tpratense_385_v2@33231
ANS_Cicer_arietinum@14208
ANS_Ahypogaea_530_v1_0@21110
ANS_Arachis_ipaensis@7786
ANS_Arachis_duranensis@9244
ANS_Lupinus_angustifolius@6416
ANS_Nelumbo_nucifera@4996
ANS_Epimedium_sagittatum@1
ANS_Aquilegia_coerulea@22072
ANS_Macleaya_cordata@5394
ANS_Setaria_italica@22738
ANS_Sviridis_500_v2_1@6441
ANS_Pvirgatum_516_v5_1@117294
ANS_Panicum_hallii@27688
ANS_Dichanthelium_oligosanthes@1451
ANS_Sorghum_bicolor@24386
ANS_Msinensis_497_v7_1@65477
ANS_Zea_mays@2
ANS_Othomaeum_386_v1_0@11759
ANS_Triticum_aestivum@1
ANS_Aegilops_tauschii@10225
ANS_Tintermedium_503_v2_1@133855
ANS_Hvulgare_462_r1@216015
ANS_Oryza_brachyantha@17888
ANS_Oryza_sativa@1
ANS_Ananas_comosus@12877
ANS_Asparagus_officinalis@19395
ANS_Allium_cepa@1
ANS_Dalata_550_v2_1@27517
ANS_Spolyrhiza_290_v2@3349
ANS_Anthurium_andraeanum@1
ANS_Zostera_marina@6565
ANS_Elaeis_guineensis@9820
ANS_Phoenix_dactylifera@22179
ANS_Musa_acuminata@11414
ANS_Tulipa_fosteriana@1
ANS_Lilium_hybrid@1
ANS_Ckanehirae_531_v3@11906
ANS_Amborella_trichopoda@11436
0.2

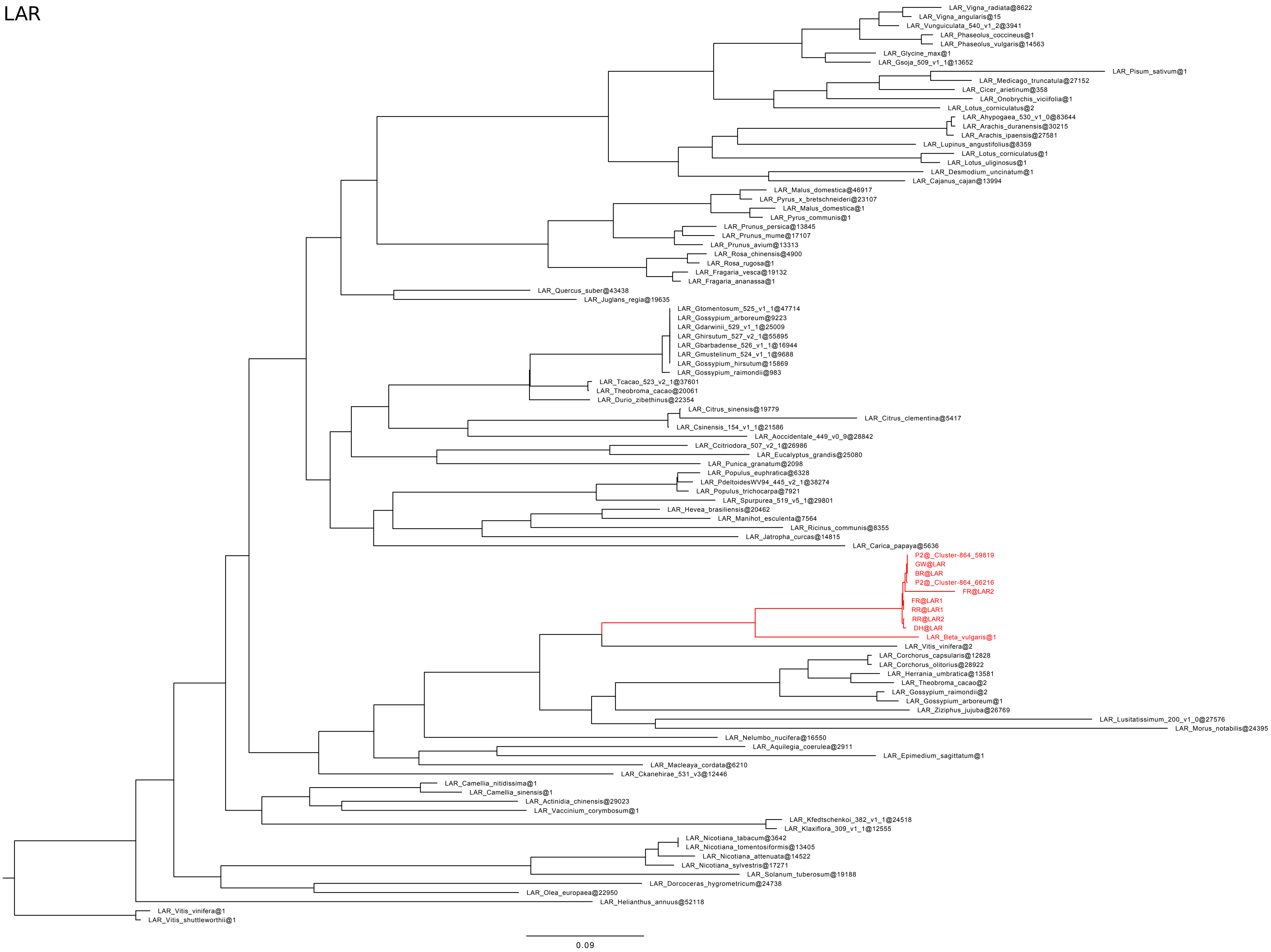
LAR
LAR_Vigna_radiata@8622
LAR_Vigna_angularis@15
LAR_Vunguiculata_540_v1_2@3941
LAR_Phaseolus_coccineus@1
LAR_Phaseolus_vulgaris@14563
LAR_Glycine_max@1
LAR_Gsoja_509_v1_1@13652
LAR_Pisum_sativum@1
LAR_Medicago_truncatula@27152
LAR_Cicer_arietinum@358
LAR_Onobrychis_viciifolia@1
LAR_Lotus_corniculatus@2
LAR_Ahypogaea_530_v1_0@83644
LAR_Arachis_duranensis@30215
LAR_Arachis_ipaensis@27581
LAR_Lupinus_angustifolius@8359
LAR_Lotus_corniculatus@1
LAR_Lotus_uliginosus@1
LAR_Desmodium_uncinatum@1
LAR_Cajanus_cajan@13994
LAR_Malus_domestica@46917
LAR_Pyrus_x_bretschneideri@23107
LAR_Malus_domestica@1
LAR_Pyrus_communis@1
LAR_Prunus_persica@13845
LAR_Prunus_mume@17107
LAR_Prunus_avium@13313
LAR_Rosa_chinensis@4900
LAR_Rosa_rugosa@1
LAR_Fragaria_vesca@19132
LAR_Fragaria_ananassa@1
LAR_Quercus_suber@43438
LAR_Juglans_regia@19635
LAR_Gtomentosum_525_v1_1@47714
LAR_Gossypium_arboreum@9223
LAR_Gdarwinii_529_v1_1@25009
LAR_Ghirsutum_527_v2_1@55895
LAR_Gbarbadense_526_v1_1@16944
LAR_Gmustelinum_524_v1_1@9688
LAR_Gossypium_hirsutum@15869
LAR_Gossypium_raimondii@983
LAR_Tcacao_523_v2_1@37601
LAR_Theobroma_cacao@20061
LAR_Durio_zibethinus@22354
LAR_Citrus_sinensis@19779
LAR_Citrus_clementina@5417
LAR_Csinensis_154_v1_1@21586
LAR_Aoccidentale_449_v0_9@28842
LAR_Ccitriodora_507_v2_1@26986
LAR_Eucalyptus_grandis@25080
LAR_Punica_granatum@2098
LAR_Populus_euphratica@6328
LAR_PdeltoidesWV94_445_v2_1@38274
LAR_Populus_trichocarpa@7921
LAR_Spurpurea_519_v5_1@29801
LAR_Hevea_brasiliensis@20462
LAR_Manihot_esculenta@7564
LAR_Ricinus_communis@8355
LAR_Jatropha_curcas@14815
LAR_Carica_papaya@5636
P2@_Cluster-864_59819
GW@LAR
BR@LAR
P2@_Cluster-864_66216
FR@LAR2
FR@LAR1
RR@LAR1
RR@LAR2
DH@LAR
LAR_Beta_vulgaris@1
LAR_Vitis_vinifera@2
LAR_Corchorus_capsularis@12828
LAR_Corchorus_olitorius@28922
LAR_Herrania_umbratica@13581
LAR_Theobroma_cacao@2
LAR_Gossypium_raimondii@2
LAR_Gossypium_arboreum@1
LAR_Ziziphus_jujuba@26769
LAR_Lusitatissimum_200_v1_0@27576
LAR_Morus_notabilis@24395
LAR_Nelumbo_nucifera@16550
LAR_Aquilegia_coerulea@2911
LAR_Epimedium_sagittatum@1
LAR_Macleaya_cordata@6210
LAR_Ckanehirae_531_v3@12446
LAR_Camellia_nitidissima@1
LAR_Camellia_sinensis@1
LAR_Actinidia_chinensis@29023
LAR_Vaccinium_corymbosum@1
LAR_Kfedtschenkoi_382_v1_1@24518
LAR_Klaxiflora_309_v1_1@12555
LAR_Nicotiana_tabacum@3642
LAR_Nicotiana_tomentosiformis@13405
LAR_Nicotiana_attenuata@14522
LAR_Nicotiana_sylvestris@17271
LAR_Solanum_tuberosum@19188
LAR_Dorcoceras_hygrometricum@24738
LAR_Olea_europaea@22950
LAR_Helianthus_annuus@52118
LAR_Vitis_vinifera@1
LAR_Vitis_shuttleworthii@1
0.09

ANR

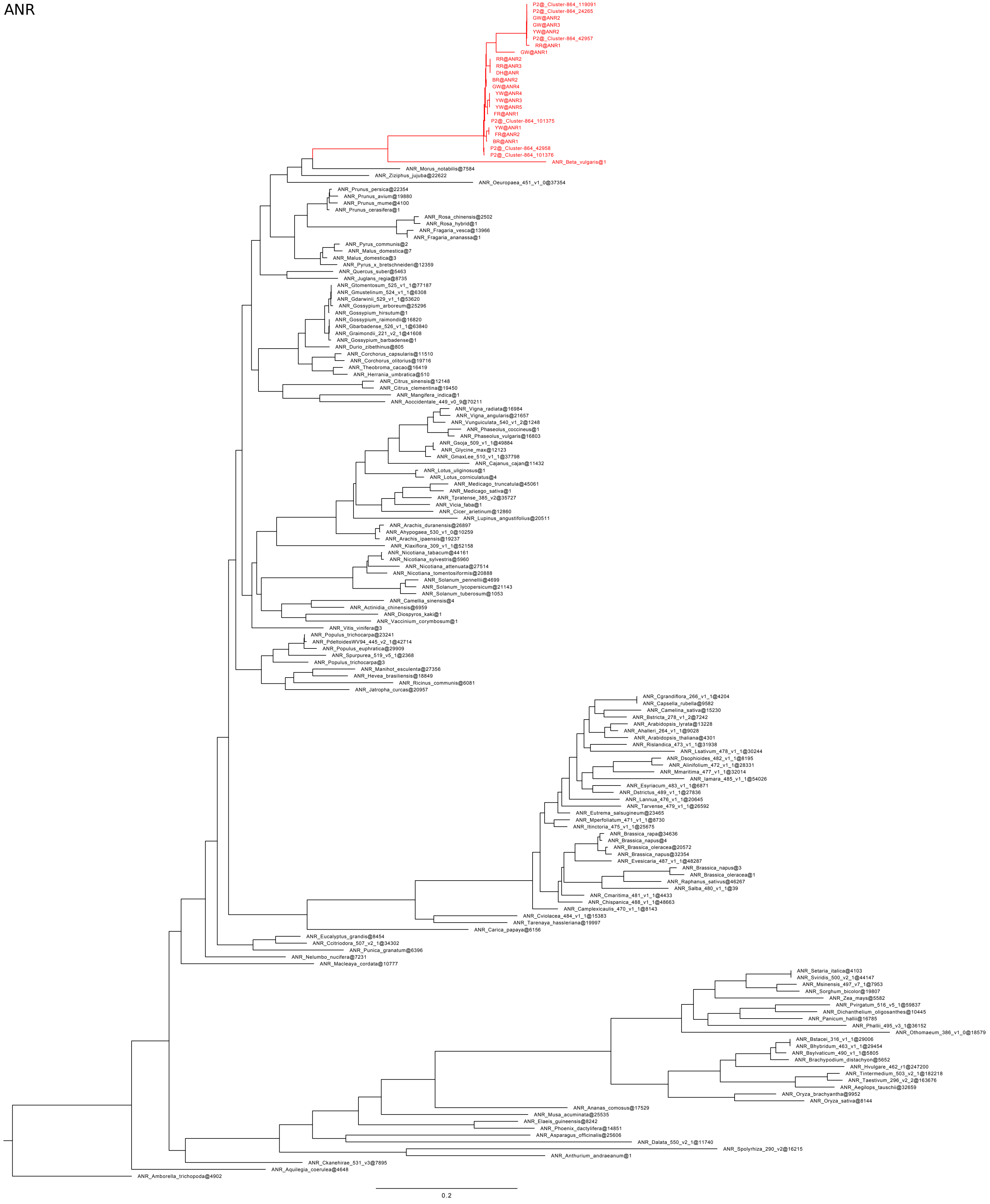
P2@_Cluster-864_119091
P2@_Cluster-864_24265
GW@ANR2
GW@ANR3
YW@ANR2
P2@_Cluster-864_42957
RR@ANR1
GW@ANR1
RR@ANR2
RR@ANR3
DH@ANR
BR@ANR2
GW@ANR4
YW@ANR4
YW@ANR3
YW@ANR5
FR@ANR1
P2@_Cluster-864_101375
YW@ANR1
FR@ANR2
BR@ANR1
P2@_Cluster-864_42958
P2@_Cluster-864_101376
ANR_Beta_vulgaris@1
ANR_Morus_notabilis@7584
ANR_Ziziphus_jujuba@22622
ANR_Oeuropaea_451_v1_0@37354
ANR_Prunus_persica@22354
ANR_Prunus_avium@19880
ANR_Prunus_mume@4100
ANR_Prunus_cerasifera@1
ANR_Rosa_chinensis@2502
ANR_Rosa_hybrid@1
ANR_Fragaria_vesca@13966
ANR_Fragaria_ananassa@1
ANR_Pyrus_communis@2
ANR_Malus_domestica@7
ANR_Malus_domestica@3
ANR_Pyrus_x_bretschneideri@12359
ANR_Quercus_suber@5463
ANR_Juglans_regia@8735
ANR_Gtomentosum_525_v1_1@77187
ANR_Gmustelinum_524_v1_1@6308
ANR_Gdarwinii_529_v1_1@53620
ANR_Gossypium_arboreum@25296
ANR_Gossypium_hirsutum@1
ANR_Gossypium_raimondii@16820
ANR_Gbarbadense_526_v1_1@63840
ANR_Graimondii_221_v2_1@41608
ANR_Gossypium_barbadense@1
ANR_Durio_zibethinus@805
ANR_Corchorus_capsularis@11510
ANR_Corchorus_olitorius@19716
ANR_Theobroma_cacao@16419
ANR_Herrania_umbratica@510
ANR_Citrus_sinensis@12148
ANR_Citrus_clementina@19450
ANR_Mangifera_indica@1
ANR_Aoccidentale_449_v0_9@70211
ANR_Vigna_radiata@16984
ANR_Vigna_angularis@21657
ANR_Vunguiculata_540_v1_2@1248
ANR_Phaseolus_coccineus@1
ANR_Phaseolus_vulgaris@16803
ANR_Gsoja_509_v1_1@49884
ANR_Glycine_max@12123
ANR_GmaxLee_510_v1_1@37798
ANR_Cajanus_cajan@11432
ANR_Lotus_uliginosus@1
ANR_Lotus_corniculatus@4
ANR_Medicago_truncatula@45061
ANR_Medicago_sativa@1
ANR_Tpratense_385_v2@35727
ANR_Vicia_faba@1
ANR_Cicer_arietinum@12860
ANR_Lupinus_angustifolius@20511
ANR_Arachis_duranensis@26897
ANR_Ahypogaea_530_v1_0@10259
ANR_Arachis_ipaensis@19237
ANR_Klaxiflora_309_v1_1@52158
ANR_Nicotiana_tabacum@44161
ANR_Nicotiana_sylvestris@5960
ANR_Nicotiana_attenuata@27514
ANR_Nicotiana_tomentosiformis@20888
ANR_Solanum_pennellii@4699
ANR_Solanum_lycopersicum@21143
ANR_Solanum_tuberosum@1053
ANR_Camellia_sinensis@4
ANR_Actinidia_chinensis@6959
ANR_Diospyros_kaki@1
ANR_Vaccinium_corymbosum@1
ANR_Vitis_vinifera@3
ANR_Populus_trichocarpa@23241
ANR_PdeltoidesWV94_445_v2_1@42714
ANR_Populus_euphratica@29909
ANR_Spurpurea_519_v5_1@2368
ANR_Populus_trichocarpa@3
ANR_Manihot_esculenta@27356
ANR_Hevea_brasiliensis@18849
ANR_Ricinus_communis@6081
ANR_Jatropha_curcas@20957
ANR_Cgrandiflora_266_v1_1@4204
ANR_Capsella_rubella@9582
ANR_Camelina_sativa@15230
ANR_Bstricta_278_v1_2@7242
ANR_Arabidopsis_lyrata@13228
ANR_Ahalleri_264_v1_1@9028
ANR_Arabidopsis_thaliana@4301
ANR_Rislandica_473_v1_1@31938
ANR_Lsativum_478_v1_1@30244
ANR_Dsophioides_482_v1_1@8195
ANR_Alinifolium_472_v1_1@28331
ANR_Mmaritima_477_v1_1@32014
ANR_Iamara_485_v1_1@54026
ANR_Esyriacum_483_v1_1@6871
ANR_Dstrictus_489_v1_1@27836
ANR_Lannua_476_v1_1@20645
ANR_Tarvense_479_v1_1@26592
ANR_Eutrema_salsugineum@23465
ANR_Mperfoliatum_471_v1_1@8730
ANR_Itinctoria_475_v1_1@25675
ANR_Brassica_rapa@34636
ANR_Brassica_napus@4
ANR_Brassica_oleracea@20572
ANR_Brassica_napus@32354
ANR_Evesicaria_487_v1_1@48287
ANR_Brassica_napus@3
ANR_Brassica_oleracea@1
ANR_Raphanus_sativus@46267
ANR_Salba_480_v1_1@39
ANR_Cmaritima_481_v1_1@4433
ANR_Chispanica_488_v1_1@48663
ANR_Camplexicaulis_470_v1_1@8143
ANR_Cviolacea_484_v1_1@15383
ANR_Tarenaya_hassleriana@19997
ANR_Carica_papaya@6156
ANR_Eucalyptus_grandis@8454
ANR_Ccitriodora_507_v2_1@34302
ANR_Punica_granatum@6396
ANR_Nelumbo_nucifera@7231
ANR_Macleaya_cordata@10777
ANR_Setaria_italica@4103
ANR_Sviridis_500_v2_1@44147
ANR_Msinensis_497_v7_1@7953
ANR_Sorghum_bicolor@19807
ANR_Zea_mays@5582
ANR_Pvirgatum_516_v5_1@59837
ANR_Dichanthelium_oligosanthes@10445
ANR_Panicum_hallii@16785
ANR_Phallii_495_v3_1@36152
ANR_Othomaeum_386_v1_0@18579
ANR_Bstacei_316_v1_1@29006
ANR_Bhybridum_463_v1_1@29454
ANR_Bsylvaticum_490_v1_1@5805
ANR_Brachypodium_distachyon@5652
ANR_Hvulgare_462_r1@247200
ANR_Tintermedium_503_v2_1@182218
ANR_Taestivum_296_v2_2@163676
ANR_Aegilops_tauschii@32659
ANR_Oryza_brachyantha@9952
ANR_Oryza_sativa@8144
ANR_Ananas_comosus@17529
ANR_Musa_acuminata@25535
ANR_Elaeis_guineensis@8242
ANR_Phoenix_dactylifera@14851
ANR_Asparagus_officinalis@25606
ANR_Dalata_550_v2_1@11740
ANR_Spolyrhiza_290_v2@16215
ANR_Anthurium_andraeanum@1
ANR_Ckanehirae_531_v3@7895
ANR_Aquilegia_coerulea@4648
ANR_Amborella_trichopoda@4902
0.2
